# The Pdgfd-Pdgfrb axis orchestrates tumor-nerve crosstalk in pancreatic cancer

**DOI:** 10.1101/2025.08.26.672505

**Authors:** Peter L. Wang, Nicole A. Lester, Ella N. Perrault, Jennifer Su, Dennis Gong, Carina Shiau, Jingyi Cao, Phuong T. T. Nguyen, Jung Woo Bae, Deniz Olgun, Hannah I. Hoffman, Ashley Lam, Jean Huang-Gao, Saifur Rahaman, Jimmy A. Guo, Jaimie L. Barth, Nicholas Caldwell, Prajan Divakar, Jason W. Reeves, Arya Bahrami, ShanShan He, Michael Patrick, Eric Miller, Maria Ganci, Grissel Cervantes Jaramillo, Theodore S. Hong, Jennifer Y. Wo, Hannah Roberts, Ralph Weissleder, Hongyoon Choi, Carlos Fernandez-del Castillo, Kathleen Cormier, David T. Ting, Tyler Jacks, Lei Zheng, Martin Hemberg, Mari Mino-Kenudson, William L. Hwang

## Abstract

Nerves are an integral component of the tumor microenvironment, contributing to cancer progression, metastasis, morbidity, and mortality. In pancreatic ductal adenocarcinoma (PDAC), worse clinical outcomes are associated with perineural invasion (PNI), a process by which cancer cells surround and invade nerves. Here, we employed whole-transcriptome and single-cell spatial transcriptomics to identify candidate tumor-nerve interactions that promote PNI. We discovered that Pdgfd signaling promotes key features of nerve invasion. Mechanistically, Pdgfd stimulated cancer cell invasiveness, neurite outgrowth, and direct physical engagement with glia. Pharmacological blockade of this axis reduced each of these processes *in vitro* as well as PNI *in vivo*. Thus, Pdgfd-Pdgfrb signaling mediates PNI by coordinating multifaceted cancer-neuron-glia interactions and represents a promising therapeutic strategy aimed at disrupting harmful cancer-nerve crosstalk.

## Introduction

Perineural invasion (PNI), a histopathologic manifestation of tumor-nerve crosstalk whereby cancer cells envelop or invade peripheral nerves, has been widely linked to adverse clinical outcomes^1–4^. One of the hallmarks of pancreatic ductal adenocarcinoma (PDAC) is an exceptionally high frequency of PNI^3,4^. Prior studies have revealed that nerve-derived signals increase the invasiveness of pancreatic cancer cells and that axonal growth and guidance factors are abundantly expressed in PDAC^5–7^. PDAC cells have also been shown to reprogram Schwann cells to promote PNI^8,9^. These observations suggest that PNI is facilitated by complex bidirectional signaling between nerves and cancer cells. However, the critical molecular mechanisms underlying these interactions are poorly understood because prior work was largely limited by a narrow focus on preselected pathways and lack of cell-type specificity and spatial context. To address this gap in knowledge, we utilized comprehensive whole-transcriptome and single-cell spatial screening approaches to identify candidate mediators of PNI from patient-derived PDAC specimens. We then performed *in vitro* and *in vivo* functional validation to establish the Pdgfd-Pdgfrb signaling axis as a multimodal mediator of PNI that facilitates cancer cell invasiveness, neurite outgrowth, and physical interactions with glial cells.

### Whole-transcriptome and single-cell resolution spatial screen for mediators of PNI

To identify pan-cancer gene expression programs associated with PNI, we previously created a custom cohort of 2,029 patients across 12 cancer types from The Cancer Genome Atlas (TCGA) with bulk transcriptomic and surgical pathology data^10^. Differential expression (DE) analysis revealed numerous genes and pathways enriched in patients with PNI, including epithelial-to-mesenchymal transition (EMT) and axonal guidance genes, as well as ones that were previously unidentified^10^. While the cellular sources and spatial context of these transcriptional programs cannot be determined from bulk data, these findings suggest that PNI is driven by cellular crosstalk that affects the programming of both cancer cells and nerves. To more comprehensively investigate tumor-nerve crosstalk and identify potential mediators of PNI, we performed a combination of whole-transcriptome digital spatial profiling (WT-DSP; Bruker/NanoString GeoMx)^11^ and 6200-plex spatial molecular imaging (6K-SMI; Bruker/NanoString CosMx) on human PDAC. This approach leverages the complementary strengths of broad molecular coverage and subcellular resolution offered by the two platforms. First, we designed twelve custom tissue microarrays (TMAs; *n* = 288 cores) derived from intratumorally-matched malignant areas, with and without nerve involvement, in resected primary PDAC specimens (*n* = 29 patients) (**Fig. S1A; Methods**). We performed WT-DSP on each of these TMAs to independently quantify transcripts from S100B^+^ nerves, αSMA^+^ cancer-associated fibroblasts (CAFs), and PanCK^+^ cancer cells within selected regions of interest (ROIs) (**Fig. 1A**). For cores with nerve involvement (N+), we defined concentric areas of interest, moving radially outward from each nerve (inner ring: <100 µm; outer ring: 100-300 µm; **Fig. S1 B and C; Methods**). For cores without nerve involvement (N-), we placed standard circular ROIs. To capture cell-type specific transcriptomic data, ROIs were partitioned by marker expression into areas of illumination (AOIs) that defined the epithelial, CAF, and nerve segments. In total, 9,401 out of 18,815 genes were detected above background, enabling the collection of high-quality whole transcriptome data from 433 ROIs (260 N+, 173 N-) (**Fig. S1B**). Next, we performed 6K-SMI on a subset of two matched TMAs containing high levels of PNI (**Fig. 1A; Methods**). We selected 43 fields of view (FOVs) across 44 cores and achieved an average transcript count of 567 per cell with 5977 genes out of 6200 detected above background.

**Figure 1.**
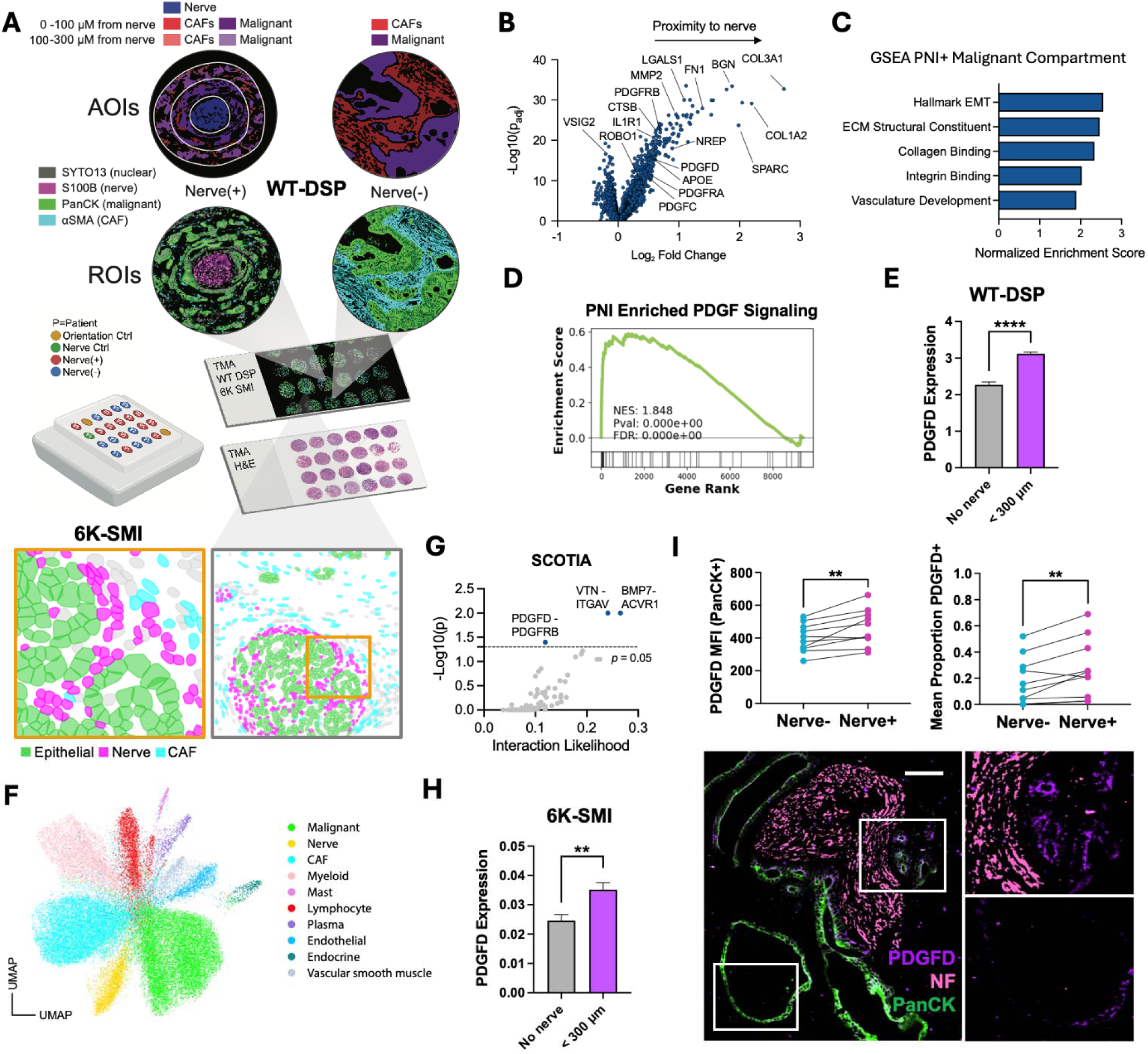
Distinct malignant-nerve signaling pathways enriched in PNI. (**A**) Schematic of spatial PNI screen with WT-DSP and 6K-SMI applied to custom PDAC TMAs. (**B**) WT-DSP volcano plot of differentially expr ssed genes in malignant cells <300 µm versus >300 µm from nearest nerve. (**C**) GSEA analysis of enriched genes expressed by malignant cells in N+ AOIs. (**D**) GSEA plot of PDGF family signaling enrichment correlated with PNI. (**E**) PDGFD expression in malignant cells located >300Lµm (no nerve, n = 126 ROIs) versus <300Lµm from the nearest nerve (n = 410 ROIs) in WT-DSP data. (**F**) UMAP of annotated cells from 6K-SMI. (**G**) SCOTIA results from 6K-SMI. (**H**) PDGFD expression in malignant cells located >300Lµm (no nerve; n = 6472 cells) versus <300Lµm from the nearest nerve (n = 6925 cells) in 6K-SMI data. (**I**) Mean fluorescence intensity (MFI) of PDGFD in PDGFD⁺ PanCK⁺ malignant cells (top left) and mean proportion of PanCK+ malignant cells that are PDGFD+ (top right) located >300Lµm (Nerve⁻) or <300Lµm (Nerve⁺) from the nearest nerve. Each dot pair represents the average of all Nerve+ and Nerve-cells analyzed in a single patient (n = 10 patients). Image (bottom left) depicts PDGFD labeling in nerve proximal (top inset) versus distal (bottom inset) PanCK+ malignant cells. Scale bar, 100 µm. *\*\*P* < 0.01, *\*\*\*\*P* < 0.0001 [Mann-Whitney test, mean SEM (E, H); paired Student’s t test, MFI and mean proportion (I)].

### Cell-type specific gene expression in PNI

We first evaluated gene expression profiles in nerves as a function of cancer cell invasion in the WT-DSP data. To facilitate this analysis, we categorized each nerve by a PNI severity score defined by the degree of malignant cell involvement (**Fig. S2A**). Higher PNI severity was associated with a progressive increase in expression of genes linked to nerve injury^12–14^, with the highest enrichment for SPP1 (**Fig. S2B**). Gene set enrichment analysis (GSEA) of PNI-associated nerves highlighted pathways related to EMT, Myc signaling, apoptosis, and autophagy (**Fig. S2B**). These observations support previous reports showing that cancer cell proximity induces nerve injury response signatures and Schwann cell reprogramming^8,15,16^. Within malignant cells, increasing severity of PNI was associated with the expression of CTDNEP1, PCBP2, and EIF4B, along with gene programs involved in Myc signaling and protein biosynthesis (**Fig. S2C**).

Next, we investigated whether transcriptional programs associated with CAF and malignant subtypes had distinct spatial associations with nerves. We scored CAFs in N+ versus N-ROIs using established subtype signatures^17^. While CAFs did not differ in inflammatory CAF (iCAF) or myofibroblastic CAF (myCAF) score as a function of nerve proximity, they were enriched in the expression of S100A11, GSTP1, and MUC1 along with gene programs associated with cytoplasmic translation, oxidative phosphorylation, and Myc signaling (**Fig. S2 D and E**). To determine whether specific malignant subtypes were associated with nerves, we scored malignant cells in N+ and N-ROIs using the subtype signatures established by Moffitt and colleagues^18^. We observed a significant enrichment of the basal-like subtype and depletion of the classical subtype in malignant cells near nerves (**Fig. S2F**). Consistent with these findings, GSEA revealed that malignant cells proximal to nerves upregulated the Hallmark EMT program (**Fig. S2G**).

To identify candidate cancer-nerve signaling pathways that may drive PNI, we focused our attention on genes enriched in malignant cells proximal to nerves. We performed differential gene expression analysis comparing malignant cells within 300 µm of a nerve versus those in N-ROIs using a linear mixed effects model (**Methods**), which yielded 1,328 enriched genes and 280 depleted genes. Among the most enriched genes were those related to extracellular matrix (ECM), including COL3A1, COL1A2, FN1, BGN, SPARC; nerve biology, including NREP, APOE, ROBO1; and cancer cell invasion, including MMP2, LGALS1, CTSB. Notably, several members of the PDGF signaling family, including PDGFD, PDGFC, PDGFRB, and PDGFRA, were also enriched (**Fig. 1B**). GSEA of the top 200 enriched genes revealed gene ontology programs related to EMT, ECM production, collagen and integrin binding, and vascular development, as well as prominent enrichment of PDGF signaling in N+ malignant cells (**Fig. 1 C and D**). Since trophic factors play a significant role in both cancer cell invasiveness and nerve recruitment^7,19^, we were interested in secreted ligands that might mediate PNI through multiple pathways. We noted that PDGFD, which was highly enriched in malignant cells near nerves (**Fig. 1E**), has previously been linked to EMT in malignant cells and neuronal development^20–22^.

To further refine and test the robustness of these results, we compared findings from the multicellular resolution WT-DSP and single-cell resolution 6K-SMI data to identify overlapping pathways. Using our previously published reference single-nucleus RNA-seq dataset^23^, we annotated the 6K-SMI data and focused on the malignant cell compartment (**Fig. 1F; Fig. S3A; Methods**). Reassuringly, genes encoding secreted ligands enriched in malignant cells near nerves were significantly correlated between the WT-DSP and 6K-SMI data (**Fig. S3B**). We used Spatially Constrained Optimal Transport Interaction Analysis (SCOTIA)^24^ to further predict paracrine ligand-receptor (LR) interactions between malignant cells and nerves. Among the statistically significant malignant-nerve LR interactions, which included BMP7-ACVR1, VTN-ITGAV, and PDGFD-PDGFRB (**Fig. 1G; Fig. S3C**), the only overlapping ligand with our DSP results was PDGFD. Indeed, malignant cell PDGFD expression in both the WT-DSP and 6K-SMI datasets showed a similar correlation with nerve proximity (**Fig. 1 E and H**). Given the identification of PDGFD signaling in PNI-associated malignant cells from both spatial transcriptomic datasets, we sought to confirm a comparable protein expression pattern by quantitative immunofluorescence (IF) imaging. Across a total of ten patients, we found significantly higher PDGFD expression in N+ versus N-malignant cells and a higher proportion of PDGFD+ malignant cells near nerves (**Fig. 1I; Fig. S4 A and B**). PDGFD expression in PanCK-negative non-malignant cells did not differ by nerve proximity (**Fig. S4C**). However, lower PanCK expression in malignant cells was correlated with PNI and nerve proximity (**Fig. S4 D and E**), consistent with a loss of epithelial identity. Collectively, these data show that malignant cells involved in PNI exhibit EMT and express distinct gene programs, including activation of the PDGFD-PDGFRB signaling axis.

### Cancer-derived Pdgfd increases nerve invasion

To investigate the potential role of Pdgfd signaling in PNI, we generated isogenic Kras^LSL-G12D/+^;Trp53^FL/FL^;Rosa26-dCas9-VPR (KP;dCas9-VPR) mouse pancreatic cancer cell lines that leverage CRISPR activation (CRISPRa) to overexpress Pdgfd or its cognate receptor Pdgfrb as well as the mNeonGreen reporter protein via *ex vivo* lentiviral transduction of Cre recombinase and guide RNA (gRNA) (**Methods**). We confirmed overexpression of Pdgfd and Pdgfrb by immunolabeling and bulk RNAseq (**Fig. 2 A and B; Fig. S5A**). We then examined programs enriched by overexpression of these genes. We found that overexpression of Pdgfd led to an increase in genes related to interferon resp nse, glycolysis, viral process, and antigen processing and presentation, and a downregulation of genes related to the regulation of epithelial cell differentiation, TGFβ signaling, membrane depolarization and regulation of cell death (**Fig. S5B**). Pdgfrb overexpression was correlated with an increase in genes related to MTORC1 signaling, synapse and cell junction organization, and regulation of cell migration, and a decrease in genes related to negative regulation of cell death, biological adhesion, vasculature development, and wound response (**Fig. S5C**).

**Figure 2.**
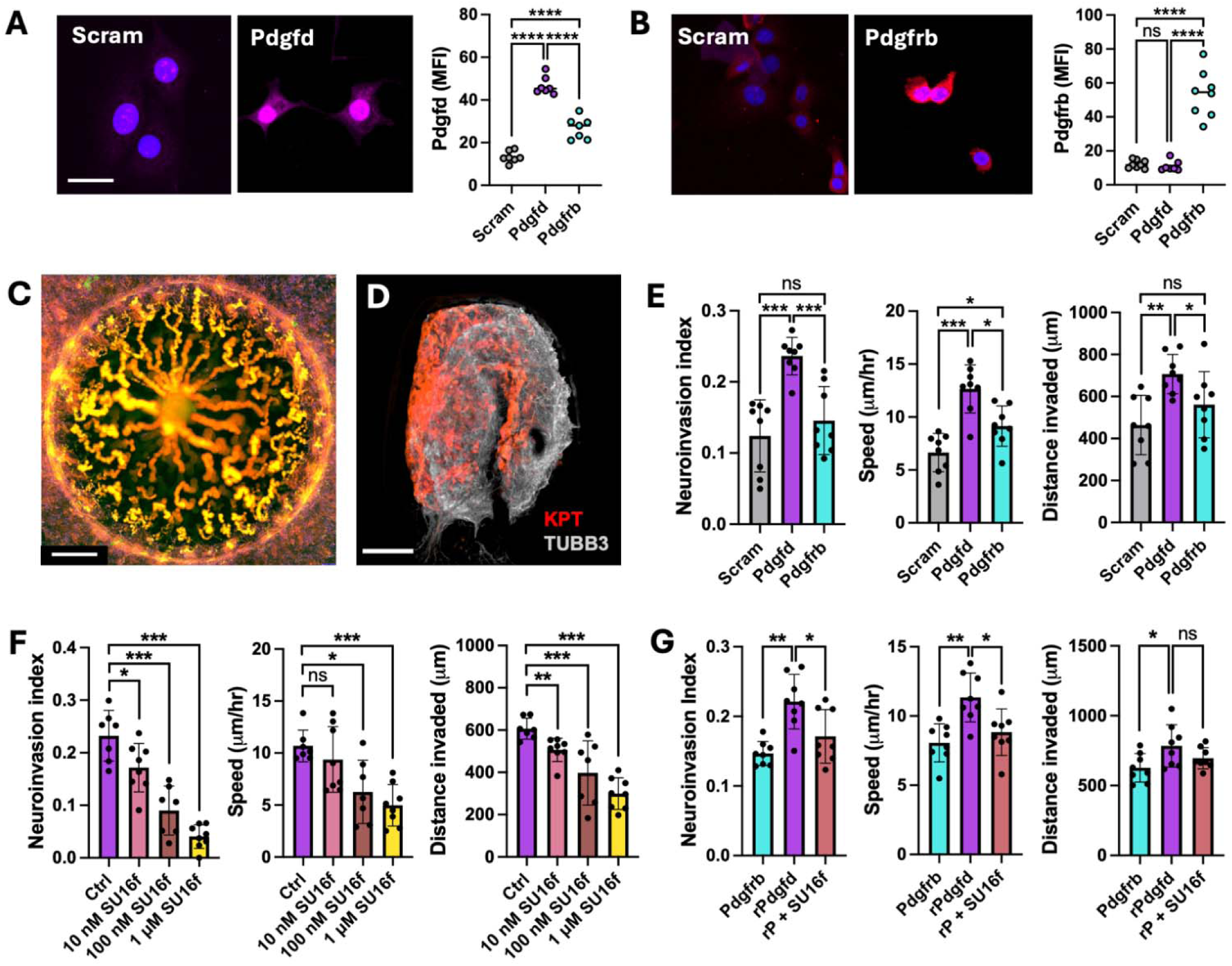
Pdgfd signaling mediates neuroinvasiveness in malignant cells. (**A, B**) MFI of Pdgfd (A) and Pdgfrb (B) in mouse PDAC cell lines expressing dCas9-VPR for CRISPR activation (n = 7-8 cells). Scale bar, 30 µm. (**C**) Whole DRG invasion assay projection image with trajectories of tdTomato+ cancer spheroids moving towards centrally located DRG over six days. Scale bar, 800 µm. (**D**) Confocal image of DRG from (C) stained with beta-3 tubulin (gray) with invaded tdTomato+ (red) cancer cells. Scale bar, 100 µm. (**E**) Quantification of neuroinvasion index, average invasion speed, and distance invaded by cancer cells upregulating candidate genes (n = 8). (**F**) Neuroinvasion index, average invasion speed, and distance traveled by Pdgfd cancer cells in co-cultures treated with varying concentrations of SU16f (n = 7-8). (**G**) Neuroinvasion index, average invasion speed, and distance traveled by Pdgfrb cells treated with vehicle, 20 nM rPdgfd, or a combination of 20 nM rP and 20nM SU16f (n = 8). *\*P* < 0.05, *\*\*P* < 0.01, \*\*\**P* < 0.001, *\*\*\*\*P* < 0.0001 [Mann-Whitney test, mean ± SD (A, B, E-G)]. Abbreviations: Scram = Scramble, rP/rPdgfd = recombinant Pdgfd.

Next, we designed a 3D neuroinvasion assay wherein a centrally-placed whole explant mouse dorsal root ganglion (DRG) embedded in a Matrigel dome is co-cultured with peripheral cancer cells to compare the neuroinvasive capacity of different cancer cell lines. We chose to use DRGs as the neuroglial source because we observed that Na_v_1.8-expressing sensory nerves^25–27^ are significantly more abundant than tyrosine hydroxylase (TH)-expressing sympathetic nerves in human PDAC (**Fig. S6A**). Cancer cells aggregate into spheroid-like structures that invade into the Matrigel dome and eventually the DRG. In contrast, cancer cell spheroids plated onto Matrigel domes without DRGs do not exhibit directed movement (**Fig. 2 C and D; Fig. S6 B-D; Movies S1 and S2**).

To quantify the neuroinvasive capacity of our isogenic cancer cell lines, we calculated the neuroinvasion index, defined as the ratio of DRG-invading cancer cell spheroids to the total number of spheroids in the Matrigel dome after six days, and measured the speed and distance traveled of invading spheroids (**Methods**). We discovered that cancer-DRG co-cultures seeded with Pdgfd cancer cells exhibited greater neuroinvasive capacity than co-cultures seeded with Pdgfrb or control (scramble gRNA) lines (**Fig. 2E; Fig. S6E**). These results suggest that expression of Pdgfd by malignant cells increases their ability to invade nerves.

Since Pdgfrb is the primary receptor for the Pdgfd homodimer^21,28^, we asked whether blocking Pdgfrb would reduce the neuroinvasiveness of Pdgfd cells. We performed the 3D neuroinvasion assay using Pdgfd cells and treated the co-cultures with SU16f, a selective and potent small-molecule inhibitor of Pdgfrb^29,30^. Su16f treatment significantly reduced the neuroinvasion index, speed, and distance traveled by Pdgfd cells (**Fig. 2F; Fig. S6F**). Next, we sought to determine whether Pdgfd is sufficient to enhance malignant cell neuroinvasion. Applying the 3D neuroinvasion assay to Pdgfrb cancer cells, we observed that supplementation with recombinant Pdgfd in the media significantly enhanced the neuroinvasion index, speed and distance traveled. These effects were reversed with the addition of SU16f (**Fig. 2g; Fig. S6G**). Our results indicate that exogenous Pdgfd enhances neuroinvasion by malignant cells.

### Pdgfd signaling increases cancer-intrinsic invasion

To determine whether Pdgfd signaling promotes malignant cell invasiveness in the absence of nerves, we performed a monoculture 2D transwell invasion assay comparing Pdgfd and control cell lines. We found that Pdgfd cells exhibited significantly higher invasiveness compared to control cells (**Fig. 3A**). To assess if the increased invasiveness of Pdgfd cells would persist *in vivo*, we performed syngeneic orthotopic transplants with 3D organoids transduced with gRNAs targeting Pdgfd or scramble control (**Methods**) as this model better recapitulates the architecture of human PDAC compared to 2D cell lines^31^. By five weeks after transplantation, Pdgfd cells invaded significantly more into the surrounding pancreas tissue compared to control cells, and mice harboring Pdgfd tumors exhibited greater pancreas mass compared to mice bearing control tumors (**Fig. 3B; Fig. S7A**). Next, we asked whether Pdgfd signaling is sufficient to increase cancer cell invasiveness. While the invasiveness of Pdgfrb was comparable to scramble control, the addition of exogenous recombinant Pdgfd induced greater invasiveness in Pdgfrb cells, though not to the extent of the Pdgfd line (**Fig. S7B**). These results demonstrate that Pdgfd increases cancer cell invasion.

**Figure 3.**
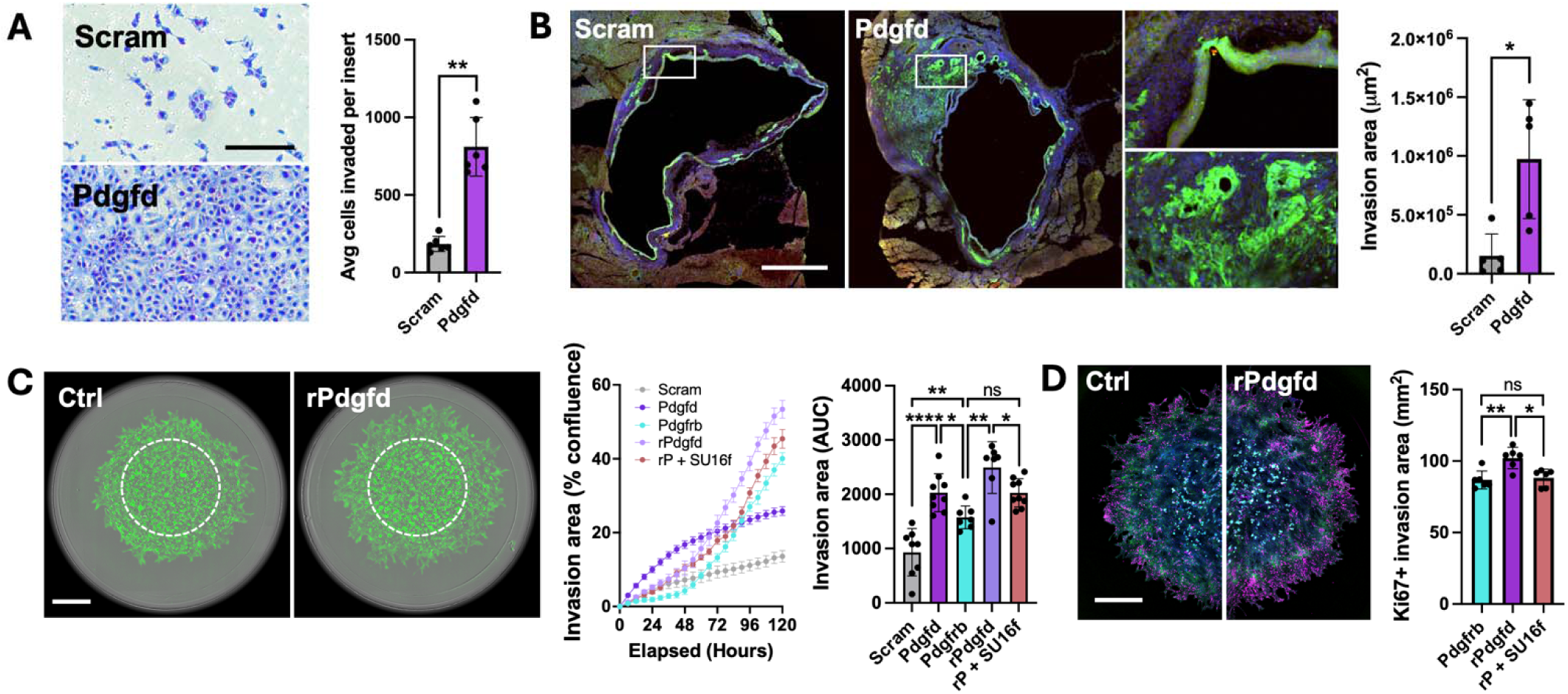
Pdgfd signaling increases cancer-intrinsic invasion. (**A**) Transwell invasion assay with scramble gRNA control and Pdgfd cells (n = 6). Scale bar, 200 µm. (**B**) Confocal imaging of 5-week orthotopically transplanted tumors (green) with scramble gRNA control and Pdgfd malignant cells (n = 5 mice). Scale bar, 800 µm. (**C**) Radial invasion assay across malignant (green) cell lines and conditions. Images show Pdgfrb (Ctrl) and Pdgfrb + 20 nM rPdgfd invasion assays at 4 days (left). Scale bar, 1 mm. Graph of invasion area over time (middle) for Pdgfrb, Pdgfrb + 20 nM rPdgfd, Pdgfrb + 20nM rPdgfd and 200nM SU16f, Pdgfd, and Scramble. Quantifi ation of AUC for invasion area (right) after 120 hours (n = 7-9). (**D**) Representative confocal image depicting invading Pdgfrb (Ctrl) and Pdgfrb + 20 nM rPdgfd cancer cells (green) stained with nuclei (blue) and Ki67 (magenta) in radial invasion assay at 5 days. Scale bar, 1 mm. Graph (right) shows Ki67+ invasion area for Pdgfrb line (Ctrl), Pdgfrb + 20 nM rPdgfd, and Pdgfrb + 20nM rPdgfd and 200nM SU16f (n = 6). *\*P* < 0.05, *\*\*P* < 0.01, *\*\*\*\*P* < 0.0001 [Mann-Whitney test, Mean SD (A-D)]. Abbreviations: Scram = Scramble, rP/rPdgfd = recombinant Pdgfd.

Since traditional 2D invasion models do not capture important characteristics of the tumor microenvironment (TME)^32,33^, we designed a 3D radial invasion assay to further assess the influence of Pdgfd on cancer cell invasiveness in a system that better reflects tumor stiffness and enables tracking over a longer period of time. Cancer cells were seeded in an inner dome of lower-density Matrigel and tracked as they invaded into an outer dome of higher-density Matrigel (**Fig. S7C; Methods**). We recapitulated the 2D invasion results, with Pdgfd cancer cells invading significantly more than Pdgfrb and scramble control lines, especially during the first 3 days. Between days 3-5, however, Pdgfrb cancer cells exhibited an accelerated invasion rate and began to surpass Pdgfd, suggesting that additional factors may drive 3D invasion dynamics over time (**Fig. S7D**). We also found that the addition of exogenous Pdgfd significantly increased the invasiveness of Pdgfrb cancer cells, even beyond that of Pdgfd cells alone, while SU16f treatment abrogated the effect of recombinant Pdgfd treatment (**Fig. 3C**). We considered whether the known mitogenic activity of Pdgfd^34,35^ might influence the invasion of Pdgfrb cancer cells. We assessed proliferation by staining for Ki67 in the radial invasion assay. Interestingly, we observed a larger area of Ki67+ cells in the recombinant Pdgfd-treated condition with proliferating cells largely confined to the invasive front. Inhibition of Pdgfd-Pdgfrb signaling with SU16f negated the increase in proliferative invading cells (**Fig. 3D**). Taken together, these results suggest that Pdgfd signaling increases cancer cell invasion through Pdgfrb and may provide a proliferative advantage to cancer cells at the invasive front challenged with a dense extracellular matrix.

### Pdgfd signaling stimulates neurite outgrowth

Since physical contact between cancer cells and nerves is an important mediator of PNI^8,19^, we hypothesized that Pdgfd may also act on nerves through neuronal Pdgfrb to drive recruitment and neurite outgrowth. First, to determine if sensory neurons express Pdgfrb, we performed flow cytometry on dissociated DRGs from Na_v_1.8-Cre Rosa26-LSL-tdTomato (Na_v_1.8-Cre-tdT) mice. We found that a subset of tdTomato+ neurons express Pdgfrb (**Fig. S8 A and B**). We further confirmed Pdgfrb expression in a subset of pancreas-innervating DRG sensory neurons in an independent dataset^36^ (**Fig. S8C**). To test whether malignant cell-derived Pdgfd promotes nerve outgrowth, we co-cultured Pdgfd, Pdgfrb, and control cancer cells with dissociated DRG sensory nerves from Na_v_1.8-Cre-tdT mice and tracked neurite outgrowth (**Fig. S8D; Methods**). We found that nerves co-cultured with Pdgfd cancer cells exhibited greater neurite outgrowth compared to those co-cultured with Pdgfrb and control cells (**Fig. 4A**). We further investigated if conditioned media (CM) from Pdgfd cancer cells was sufficient to induce the same effect. Indeed, neurons grown in Pdgfd cancer cell CM exhibited greater neurite outgrowth compared to neurons grown in control media (**Fig. 4B**). Because CM may contain additional growth factors, we sought to determine whether Pdgfd CM-enhanced neurite outgrowth occurs specifically through Pdgfrb signaling. We blocked Pdgfrb with SU16f in Pdgfd CM-treated neurons and found that neurite outgrowth was attenuated (**Fig. 4B**). To determine whether Pdgfd alone is sufficient to increase neurite outgrowth, we treated neurons with recombinant Pdgfd and observed a concentration-dependent increase (**Fig. 4C**). Finally, since intrapancreatic ganglia are a major source of pancreatic innervation, we also tested these conditions using whole DRG explants embedded in Matrigel. We observed that recombinant Pdgfd increased whole DRG neurite outgrowth, which could be prevented by SU16f treatment (**Fig. 4D**). Taken together, these results indicate that Pdgfd signaling is sufficient to induce neurite outgrowth.

**Figure 4.**
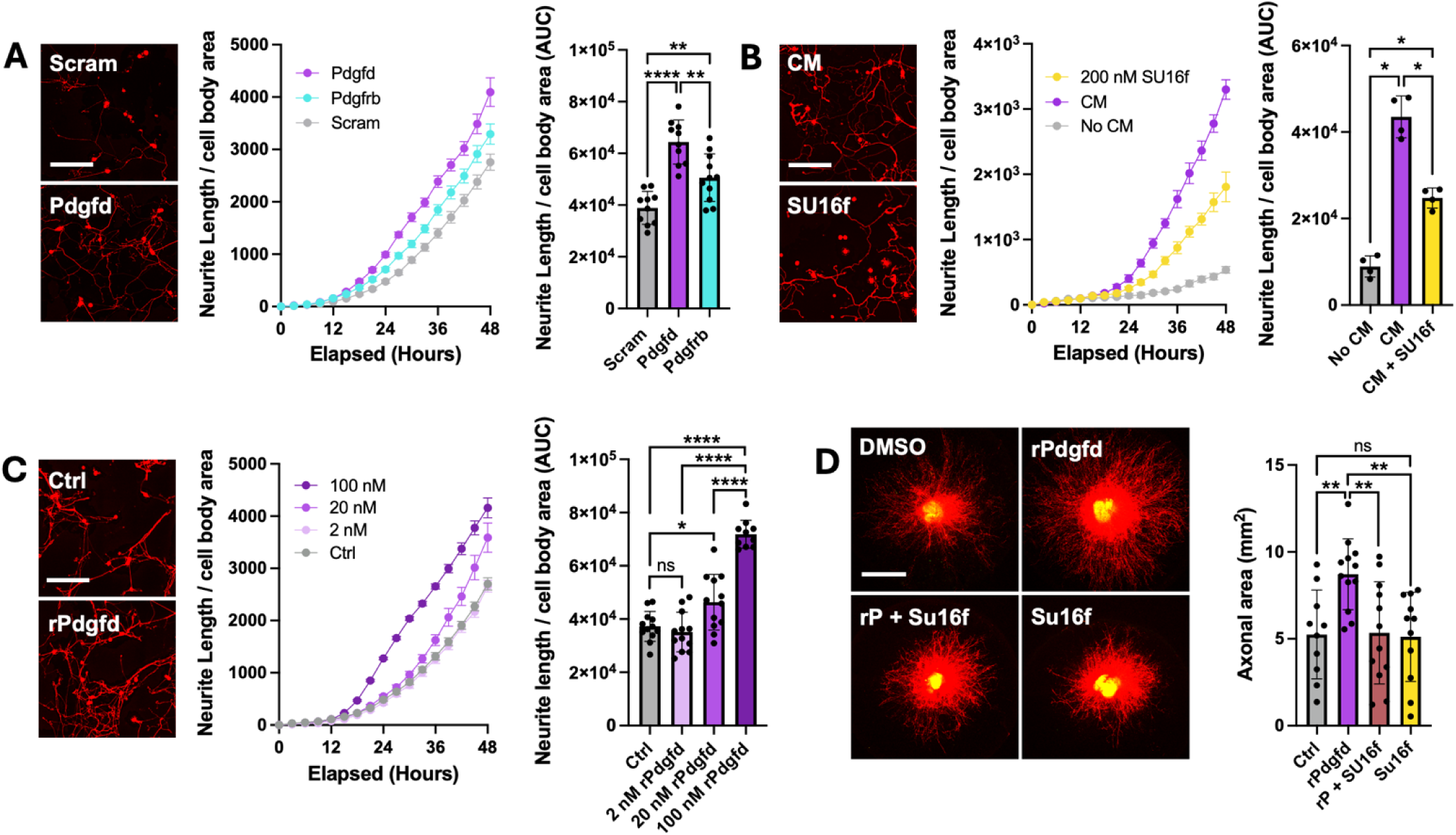
Pdgfd signaling enhances nerve outgrowth. (**A**) Representative images and growth curves of nerves from Na_v_1.8-Cre tdTomato+ mice co-cultured with Pdgfd, Pdgfrb, or scramble gRNA control cancer cells (left). Scale bar, 200 µm. Quantification (right) of area under the curve (AUC) for neurite growth curves after 48 hours (n = 10-11). (**B)** Images and growth curves of nerves grown in control media, treated with conditioned media (CM) from Pdgfd-overexpressing malignant cells, and treated with a combination of CM and 200 nM of Su16f (left). Scale bar, 200 µm. Quantification (right) of AUC for neurite growth curves after 48 hours (n = 4). (**C**) Images and growth curves of nerves treated with control vehicle, 2 nM, 20 nM, and 200 nM of rPdgfd (left). Scale bar, 200 µm. Quantification (right) of AUC for neurite growth curves after 48 hours. (**D**) Images and quantification of axonal area in DRGs treated with vehicle (Ctrl), 20 nM rPdgfd, 20 nM rPdgfd and 200 nM SU16f, and 200 nM SU16f (n = 10-12). Scale bar, 1 mm. *\*P* < 0.05, *\*\*P* < 0.01, *\*\*\*\*P* < 0.0001 [Mann-Whitney test, Mean SD (A-D)]. Abbreviations: Scram = Scramble, rP/rPdgfd = recombinant Pdgfd.

### Pdgfd-Pdgfrb pathway mediates cancer-glial interactions

Pdgfrb signaling facilitates neuronal development and growth through glial precursors and is involved in the injury response of glial cells, including Schwann cells and migratory neural crest-derived mesenchymal cells^37–40^. Since reprogrammed Schwann cells have also been shown to promote nerve invasion in pancreatic cancer^8,9^, we hypothesized that cancer cell-derived Pdgfd may drive PNI by signaling to Pdgfrb-expressing glial cells. First, we sought to confirm Pdgfrb expression in glial cells by performing flow cytometry and fluorescence imaging on primary DRGs isolated from Plp-EGFP mice, which express an enhanced green fluorescent protein (EGFP) in glial cells such as Schwann cells and their progenitors^41–43^. In freshly dissociated DRGs, we found that a subpopulation (29.6%) of EGFP+ glia expressed Pdgfrb (**Fig. S8E**). Interestingly, after culturing primary DRG cells for four days, the proportion of Pdgfrb+ glia shifted significantly to nearly all (98.7%) EGFP+ cells (**Fig. 5A; Fig. S8F**). These results show that while Pdgfrb is expressed in a subset of glial cells in naive DRGs, its expression can be increased after *ex vivo* manipulation, suggesting sensitivity to tissue perturbation.

**Figure 5.**
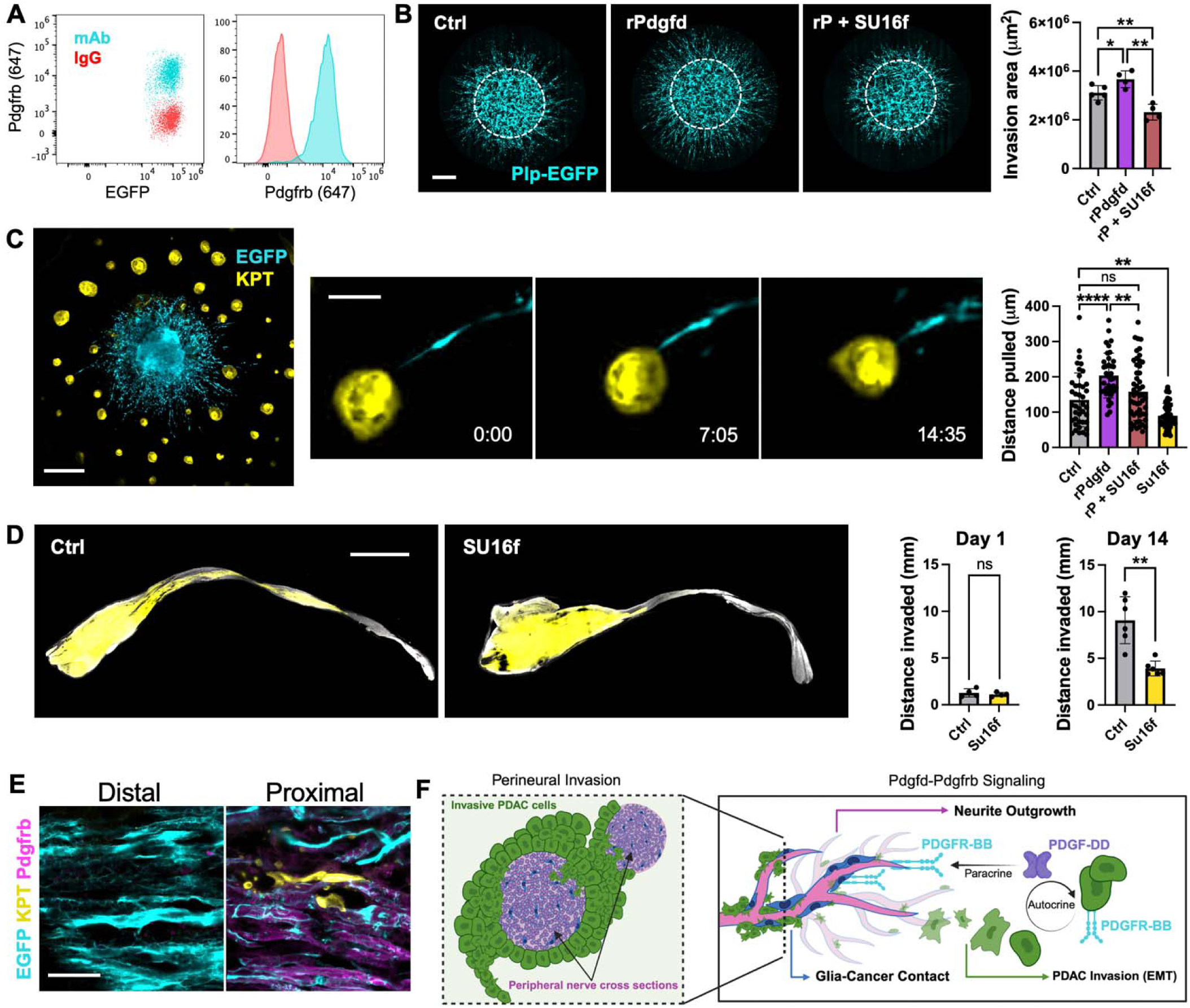
Pdgfd-Pdgfrb signaling between cancer cells and glial cells. (**A**) Flow plot and histogram showing Pdgfrb expression in EGFP+ glial cells cultured for 4 days. (**B**) Radial invasion assay with dissociated EGFP+ glial cells (left). Cultures were treated with vehicle (Ctrl), 20 nM rPdgfd, or 20 nM rPdgfd and 20 nM SU16f. Scale bar, 1 mm. Quantification (right) of invasion area at 96 hours (n=4). (**C**) Live confocal image of DRG invasion with Plp-EGFP+ DRG (cyan) and KPT cancer cells (yellow; left). Scale bar, 500 µm. Still frames with elapsed time in hours and minutes of an EGFP+ glial cell (cyan) pulling a cancer spheroid (yellow) towards the DRG (middle). Scale bar, 100 µm. Analysis of distance tdTomato+ cancer spheroids were pulled by EGFP+ glia over 24 hours (n = 37-51 spheroids per group). (**D**) Images of sciatic nerves injected with KPT cancer cells from mice treated daily with vehicle (Ctrl) or SU16f daily for 14 days (left). Scale bar, 2 mm. Quantification of sciatic invasion distance (right) at days 1 and 14 (n = 4-6 mice). (**E**) Sciatic nerve cross-section from Plp-EGFP mouse injected with KPT cancer cells. Pdgfrb (magenta) is absent in glial cells distal to invading cancer cells (left) and present in glial cells proximal to tdTomato+ cancer cells (right). Scale bar, 40 µm. (**F**) Schematic of proposed Pdgfd-Pdgfrb signaling model. *\*P* < 0.05, *\*\*P* < 0.01, *\*\*\*\*P* < 0.0001 [Unpaired t test, Mean SD (B); Mann-Whitney test, Mean SD (C, D)]. Abbreviations: rP/rPdgfd = recombinant Pdgfd.

To directly assess whether Pdgfrb expression in glia is involved in mediating the neural response to Pdgfd, we performed the radial invasion assay using glial cells. Specifically, we seeded dissociated Plp-EGFP+ glial cells in lower-density inner Matrigel domes and tracked their outward migration into surrounding higher-density Matrigel. When we added exogenous Pdgfd to this assay, we saw a significant increase in the expansion and radial migration of EGFP+ glia, which was reversed with Su16f treatment (**Fig. 5B**). To further investigate whether EGFP+ glia interact with cancer cells, we performed live confocal imaging on co-cultures of mouse Plp-EGFP DRGs and *Kras*^LSL-G12D/+^;*Trp53*^FL/FL^;tdTomato (KPT) cancer cells (**Fig. 5C; Methods**). Remarkably, we observed pulling of KPT cancer cells along nerves by EGFP+ glia towards the DRG neuronal cell bodies (**Fig. 5C**; **Movie S3**). Using a 3D phase holotomography imaging approach, we further validated the phenomenon of cancer cells being pulled along nerves (**Movie S4**). These observations are consistent with prior findings on cancer-glial interactions in PDAC models, wherein cancer-associated Schwann cells physically push and pull cancer cells to modulate their movement along nerves^8,9^. We found that the addition of recombinant Pdgfd to the whole DRG co-culture increased the distance that cancer cells were pulled by EGFP+ glia, while Su16f decreased the pulling distance (**Fig. 5C**). Taken together, these data show that Pdgfd signaling increases glial cell mobilization and that cancer-glia interactions are mediated through the Pdgfd-Pdgfrb axis.

### Inhibiting Pdgfd-Pdgfrb axis reduces PNI *in vivo*

To assess the effects of Pdgfd-Pdgfrb signaling on nerve invasion *in vivo* in a system that contains both axons and glial cells, we adopted a sciatic nerve invasion model in which cancer cells are injected into a mouse sciatic nerve and allowed to invade proximally towards the spinal cord (**Methods**). We injected KPT cells into sciatic nerves of wild-type C57BL/6J mice for quantification of invasion with or without SU16f treatment. After 14 days, mice that received daily intraperitoneal (i.p.) injections of SU16f exhibited significantly decreased nerve invasion compared to vehicle control (**Fig. 5D**). To further visualize cancer-glial interactions, we injected KPT cells into sciatic nerves of Plp-EGFP mice and found that Pdgfrb expression was increased in EGFP+ glial cells proximal, but not distal to invading KPT cells (**Fig. 5E**). These results suggest that cancer invasion of nerves causes tissue perturbations that trigger reprogramming and Pdgfrb upregulation in Schwann cells. Collectively, our data supports a previously unknown mechanism facilitating PNI in which Pdgfd-Pdgfrb signaling has multimodal effects among cancer cells, neurons, and glial cells to increase cancer cell invasiveness, nerve outgrowth, and glial cell migration and physical interactions with cancer cells (**Fig. 5F**).

## Discussion

Cancer cell crosstalk with peripheral nerves is an emerging area of interest with significant implications for disease progression and metastasis, tumor immunity, and patient quality of life^1–4,44^. While previous studies have been limited to unidirectional signaling, preselected pathways, and a lack of spatial context, our study utilized a comprehensive whole-transcriptome and single-cell spatially-resolved screen of human PDAC to identify candidate mediators of PNI. We discovered that the growth factor Pdgfd, expressed by cancer cells in the TME, drives PNI by increasing cancer cell invasiveness as well as nerve recruitment and outgrowth. Conceptually, our work reveals for the first time how a single conserved signaling axis can influence multiple avenues of tumor-nerve crosstalk across cancer cells, neurons, and glia. More broadly, this study demonstrates the meaningful integration of spatial transcriptomics and functional biological interrogation to elucidate impactful new biology.

The PDGF/PDGFR signaling axis plays essential roles in organogenesis, tissue regeneration, and wound healing, particularly in the development and function of mesenchymal cells ^22,45^. As cancer mimics a non-resolving wound^46–48^, PDGFD–PDGFRB signaling may be aberrantly activated in an attempt to carry out its physiological role in promoting EMT. In light of recent studies which link EMT cancer cell states to nerve proximity^10,49,50^, our observations suggest that PDGFD signaling may coordinate regenerative programs within the TME involving both cancer–cancer and cancer–nerve interactions.

While nerves are increasingly recognized as a central part of the TME, they have long been known to be crucial for tissue regeneration^51–53^. Importantly, the neural contribution to wound repair is strongly linked to glial cells, which employ regenerative signaling pathways similar to those of EMT^37,54,55^. In particular, their response to tissue injury has been shown to involve the Pdgfa-Pdgfra signaling axis that glial precursor cells use to migrate into wound beds and promote tissue regeneration^37,51,56^. Moreover, Pdgfa and Pdgfb have been shown to initiate neurite outgrowth^57^. While the Pdgfd-Pdgfrb axis is less recognized in non-malignant nerve injury responses, our work shows that Pdgfd signaling induces nerve outgrowth and glial mobilization in the cancer context. From this perspective, cancer cell-derived Pdgfd may recruit nerves as part of a maladaptive wound repair response in the TME. Furthermore, while previous work has linked PNI-induced nerve injury with Schwann cell dedifferentiation and subsequent involvement in progressing nerve invasion^8,54,58^, the mechanisms governing cancer-glial interactions remained unknown. Our work identifies Pdgfd-Pdgfrb signaling as one such pathway that can be targeted.

In summary, this study harnessed the cell-type specificity, spatial resolution, and comprehensive molecular coverage of a combined spatial transcriptomic screen on human PDAC to identify potential mediators of PNI. While we were limited in the number of candidate interactions that we could functionally investigate, our data converged on the PDGFD-PDGFRB axis as an orchestrator of processes spanning cancer cell invasion, neurite outgrowth and recruitment, and contact-dependent glial interactions to facilitate perineural invasion. Targeting PDGFD-PDGFRB signaling may represent an effective therapeutic strategy against cancer-nerve crosstalk for improving patient outcomes.

## Materials and methods

### Ethics

All patients in this study consented without compensation to the excess tissue biobank protocol 2003P001289 (principal investigator: C.F.C.; co-investigator: M.M.K., W.L.H.) and samples were obtained through the secondary use protocol 2022P001264 (principal investigator: W.L.H.), which was reviewed and approved by the Massachusetts General Hospital Institutional Review Board. This study is compliant with all relevant ethical regulations.

### Human tumor specimens

Two independent patient cohorts were utilized in this study. All patients included in this study had upfront resectable, nonmetastatic pancreatic ductal adenocarcinoma (PDAC) and were treatment naïve. Board-certified pathologists specializing in gastrointestinal malignancies (M.M-K., N.C.) identified regions of interest and specific pathological features within formalin-fixed paraffin-embedded (FFPE) tissue blocks. For the first cohort, we created twelve custom tissue microarrays (n=288 cores, 3 mm diameter) derived from intratumorally-matched regions with and without perineural invasion (PNI) in primary PDAC specimens (n=29 patients) to be used for whole-transcriptome digital spatial profiling (Bruker Spatial Biology/Nanostring). In the second cohort, regions containing cancer and instances of PNI were identified in primary PDAC specimens (n=10). These FFPE blocks were used for consecutive section multiplex immunostaining, fluorescence imaging, and quantification.

### Animals

All animal studies described in this study were approved by the MIT and MGH Institutional Animal Care and Use Committees (IACUC). Generation of *Kras*^LSL-G12D/+^; *Trp53*^FL/FL^ (KP) mice has previously been described^59,60^ and were bred in-house. Generation of KP;Rosa26-CAG-LSL-dCas9-VPR-P2A-mNeonGreen mice has previously been described^61^ and were bred in-house. To generate Na_v_1.8-Cre-tdTomato mice, homozygous Ai14 mice (JAX, 007914) were crossed with homozygous Na_v_1.8-Cre mice (JAX, 036564). Plp-EGFP mice^41^ were generously gifted by Allan M. Goldstein.

### Whole-transcriptome digital spatial profiling (WT-DSP)

We followed published experimental methods (Bruker Spatial Biology/Nanostring) with slight modifications^23^. FFPE PDAC specimens were examined by a pathologist, and selected cores were arranged into 12 TMAs. Serially sectioned (5µm) TMAs were processed for H&E and whole transcriptome digital spatial profiling (WT-DSP), respectively. For WT-DSP, histopathologic guidance from a consecutive hematoxylin and eosin (H&E) section was used to identify N+ and N-cores. Slides were baked for 11 hours at 37°C and 1 hour at 60°C, then promptly deparaffinized in CitriSolv (Decon, 1601H), rehydrated, antigen-retrieved in 1x Tris-EDTA/pH 9 in a steamer for 15 minutes at 10°C, digested in 0.1 ng/mL proteinase K (Thermo Fisher Scientific, AM2548) for 15 minutes at 37°C, postfixed in 10% neutral-buffered formalin for 10 minutes, hybridized to UV-photocleavable barcode-conjugated RNA in situ hybridization probe set for 24 hours at 37°C, and washed to remove unbound probes. The slides are then blocked and counterstained with morphology markers at room temperature in Buffer W (Bruker Spatial Biology/Nanostring) for 1 hour each.

The morphology markers included: SYTO13 (Thermo Fisher Scientific, S7575), anti-PanCK-Alexa Fluor 532 (Novus Biologicals, NBP2-33200AF532), anti-S100B-Alexa Fluor 594 (Novus Biologicals, NBP2-54426AF594), anti-αSMA-Alexa Fluor 647 (R&D Systems, IC1420R-100UG), all used at a 1:100 dilution. These four markers permitted delineation of nuclear, epithelial, nerve, and fibroblast compartments, with exposure set to 150 m/s, 300 m/s, 150 m/s, and 150 m/s, respectively, on the GeoMx DSP Instrument (DSP). ROIs (600 µm diameter) were selected based on the immunofluorescence image, and PNI negative ROIs were segmented directly on the instrument to create epithelial and fibroblast AOIs. PNI positive ROIs were downloaded and segmented separately on ImageJ FIJI with a custom script. The custom script created AOIs for the nerve, as well as epithelial and fibroblasts within 100 μm and 300 μm from the nerve. Library preparation was performed according to the manufacturer’s protocol (Bruker Spatial Technologies). The pooled libraries were sequenced using NovaSeq 6000 S2 flow cells following the specifications provided by Bruker Spatial Technologies (paired-end reads at length 27, index length 8) with 5% PhiX spike-in.

### WT-DSP data preprocessing and quality control

We used the GeoMx NGS Pipeline software (v2.0) to process Illumina FASTQ sequencing files to GeoMx readable digital counts (DCC). We loaded the DCC, PKC, and annotation files to R (v4.1.2) for further processing using the GeoMxTools (v2.0.0) gene expression analysis workflow. We used the PKC (probe assay metadata) file named Hs_R_NGS_WTA_v1.0.pkc, downloaded from NanoString Technologies. The annotation file contains information on the sample name, slide name, ROI number, segment type, and segment area for every collected AOI. Before proceeding with quality control and preprocessing, we first shifted all gene expression counts with a value of 0 to 1 to better enable downstream transformations.

We assessed the quality of each AOI segment. We removed AOI segments with more than 1000 raw sequencing reads; less than 80% of aligned, stitched, or trimmed sequencing reads; less than 50% sequencing saturation ([1-deduplicated reads/aligned reads]%); less than 2 negative control counts; more than 3000 counts observed in the non-template control (NTC) well; and less than 1000 in segment area. We also removed AOI segments with negative control probe expression that had a geometric mean of less than 2.

We then removed low-performing negative control probes from specific segments) or globally (across the entire study). Locally, we removed negative control probes that were determined as outliers according to the Grubbs test. Globally, we removed negative control probes that either had a geometric mean from all segments divided by the geometric mean of all negative probe counts less than 0.1, or were an outlier (according to the Grubb’s test) in at least 20% of segments.

We determined the limit of quantification (LOQ) per segment. We used two geometric standard deviations above the geometric mean as the LOQ threshold. We also designated a minimum LOQ of 2 to be used if the LOQ calculated in a segment fell below this threshold. We filtered segments with exceptionally low signal, namely those with less than 5% of panel genes detected above the LOQ relative to other segments. We only focused on genes detected in at least 5% of segments.

### WT-DSP normalization and detrending

We normalized the data using upper quartile (Q3) normalization. We then detrended the log-transformed normalized expression by first calculating adjustment factors, which have been adapted from our previously published work :

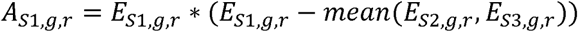

Adjustment factors *A* are calculated for the expression *E* of gene *g* in ROI *r* by comparing a given segment *S1* to the average expression in adjacent segments, *S2* and *S3*. The original expression of segment *S1* is then detrended using the following:

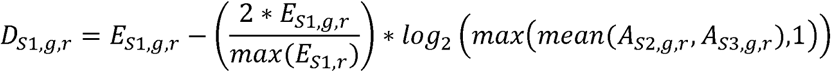

For each AOI, there may be one, two, or three adjacent segments, which necessarily modifies the formulas above. For N+ ROIs, nerve AOIs (*S1*) were detrended against epithelial (*S2*) and CAF (*S3*) AOIs in the inner ring; epithelial AOIs (*S1*) in the inner ring were detrended against the nerve AOI (*S2*) and CAF AOIs (*S3, S4*) in the inner and outer rings; CAF AOIs (*S1*) in the inner ring were detrended against the nerve AOI (*S2*) and epithelial AOIs (*S3, S4*) in the inner and outer rings; epithelial AOIs (*S1*) in the outer ring were detrended against CAF AOIs (*S2, S3*) in the inner and outer rings; and CAF AOIs (*S1*) in the outer ring were detrended against epithelial AOIs (*S2, S3*) in the inner and outer rings. However, we made an exception for N+ ROIs with only neurites. For these ROI, both epithelial AOIs (*S1*) are detrended against CAF AOIs (*S2, S3*) in the inner and outer rings, but not against the nerve AOI. Similarly, both CAF AOIs (*S1*) are detrended against epithelial AOIs (*S2, S3*) in the inner and outer rings. For these ROI, the treatment of nerve AOIs is the same. For N-ROIs, epithelial AOIs (*S1*) are detrended against CAF AOIs (*S2*), and CAF AOIs (*S1*) are detrended against epithelial AOIs (*S2*).

Since a substantial number of AOIs have been filtered via quality control, some N+ ROIs do not have data collected for all segments. To correct for the impact that a missing AOI *m* will have on the detrended expression of an adjacent AOI *a*, we first perform K-means clustering on the subset of AOIs of the same cell type as AOI *a* that additionally have recorded expression values for an adjacent AOI of the same cell type as AOI *m*. We then computed a placeholder expression value for the missing AOI *m* by calculating the mean expression of the adjacent AOIs of the same cell type as AOI *m* that are in the same clustering assignment. Placeholder expression values are used in detrending computations.

### WT-DSP analysis on nerve proximity and snRNA-seq programs

To compute the normalized program scores in each ROI, we first calculated the mean expression of the top 50 genes from snRNA seq-derived malignant and CAF lineage programs described in our prior work^23^. We performed L1 normalization on the program scores such that the scores sum to 1 in each ROI. We then scaled all of the program scores, in aggregate, using min-max normalization. This normalization is done separately for malignant (MES, CLS) and CAF (ADH, IMM, NRT, MYO) lineage programs. To compare the normalized program expressions in malignant or CAF segments within N+ *versus* N-ROIs, we used a two-sided Mann-Whitney test with Bonferroni-correction. For some analysis, we further stratified N+ ROIs into those with large nerves and those with small neurites, and N+ ROIs with nerves neither classified as large nor small are removed.

### WT-DSP differential gene expression analysis

We performed differential expression (DE) analysis using a linear mixed-effects model (LMM; lme4 v1.1-31 and lmerTest v3.1-3) to compare two populations of AOIs to identify enriched and depleted genes, in association with nerve proximity:

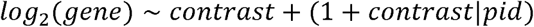

where the random slope is the contrast (e.g., the variable distinguishes the two populations of AOIs, such as N+ *versus* N-) and the random intercept is the patient to which the AOI belonged. We applied a significance threshold using a false discovery rate (FDR) equal to 0.05.

### 6K spatial molecular imaging workflow

Tissue samples were prepared for spatial molecular imaging on a pre-commercial SMI platform (Bruker Spatial Biology/Nanostring) following the manufacturer’s instructions. Slides with TMA sections were baked overnight at 60°C to enhance FFPE tissue adherence and were processed following the CosMx RNA assay manual slide preparation manual with minor modifications. Briefly, samples underwent deparaffinization, proteinase K (1 μg/ml; ThermoFisher) digestion at 40°C for 30 minutes, and heat-induced epitope retrieval (HIER) at 100°C for 15 minutes in CosMx target retrieval buffer to expose target RNAs and epitopes. The samples were rinsed with diethyl pyrocarbonate (DEPC)-treated water (DEPC H2O) twice before incubating with 1:1000 diluted fiducials (Bangs Laboratory) in 2X saline sodium citrate, 0.001% Tween 20 (SSCT) solution for 5 minutes. The samples were rinsed with 1X PBS to remove excess fiducials and fixed with 10% neutral buffered formalin for 5 minutes. Fixed samples were washed with Tris-glycine buffer (0.1M glycine, 0.1M Tris-base in DEPC H2O) and 1X PBS for 5 minutes each before being blocked using 100 mM N-succinimidyl acetate (NHS-acetate, ThermoFisher) in CosMx NHS-acetate buffer for 15 minutes. Prepared samples were rinsed with 2X saline sodium citrate for 5 minutes, after which an Adhesive SecureSeal Hybridization Chamber (Grace Bio-Labs) was placed on the slide to cover the samples. RNA ISH probes (CosMx 6000-plex probe mix) were denatured at 95°C for 2 minutes and then placed on ice before preparing the ISH probe mix (1 nM ISH probes, 1X Buffer R, 0.1 U/μl SUPERaseIn™ in DEPC H2O). The ISH probe mix was pipetted into the hybridization chamber, and hybridization was performed at 37°C overnight. After overnight hybridization, samples were washed twice with 50% formamide in 2X SSC at 37°C for 25 minutes, rinsed twice with 2X SSC for 2 minutes. The DAPI nuclear stain was applied to the samples at 5 µg/mL for 15 minutes at room temperature. Following nuclear staining samples were washed in PBS for 5 min, the CosMx RNA assay nuclear (DAPI) and cell segmentation antibody mix containing CD298/B2M (Abcam, EP1845Y)/(Abcam, EP2978Y), PanCK (Novus, AE-1/AE-3), CD45 (Novus, 2B11 + PD7/26) and CD68 (Novus, C68/684) was applied in RNA Blocking Buffer (Bruker Spatial Biology) and incubated with the sample at room temperature for 2 hours. Slides were then washed three times in PBS for 5 minutes each, and a CosMx flow cell was applied prior to loading onto the CosMx instrument. Fields of view (FOVs), each measuring 0.51 mm x 0.51 mm, were selected and target RNA readout on the SMI instrument was performed, following published protocols^62^. After the CosMx instrument run was completed, a custom on-instrument workflow was applied to stain the tissue with AF647 conjugated S100B (Abcam, 215989) and B3 Tubulin (Abcam, ab221935) antibodies.

### 6K-SMI image processing and cell segmentation

Raw image processing and feature extraction were performed using an in-house SMI data processing pipeline, which includes registration, feature detection, and localization. The Z-stack images of nuclear (DAPI) and CD298/B2M (surface) staining were used for drawing cell boundaries on the samples. A cell segmentation pipeline using *Cellpose* (*v2.0.5*)^63,64^ was used to accurately assign transcripts to cell locations and subcellular compartments. Segmentation robustness was evaluated by comparing the results of another cell segmentation tool *Baysor* (*v0.6.0*)^65^. The transcript profile of individual cells was generated by combining target transcript location and cell segmentation boundaries. Cells with fewer than 60 total transcripts were omitted from downstream analysis. Expression profiles were normalized for each cell by dividing its raw count vector by total counts and multiplying by the scale factor of 10,000 before log transformation.

### 6K-SMI data analysis

Raw image processing and cell segmentation were performed using the strategy we previously described^24^. Cells with fewer than 100 total transcripts were omitted from downstream analysis. Cell-type identification was conducted using Seurat (v4.2.0) in R. Gene expression profiles were normalized for each cell by dividing its raw count vector by total counts, multiplying by a scale factor 10,000 before log transformation. Highly variable genes were identified using the FindVariableFeatures function. Principal component analysis (PCA) was performed on the scaled expression data of highly variable genes. To correct for potential batch effects, Harmony integration was applied using the RunHarmony function. Clustering was performed using the NBclust (v3.0.1) to determine the optimal number of clusters. Cell clusters were visualized using Uniform Manifold Approximation and Projection (UMAP) dimensionality reduction. Differentially expressed genes (DEGs) for each cluster were identified using the FindMarkers function with the Wilcoxon rank-sum test.

### 6K-SMI cell type annotations

Cell type annotation was performed using *Insitutype* (*v1.0.0*)^66^ for supervised clustering. High-confidence malignant and non-malignant cells were selected as anchors based on protein (PanCK) and expression (*KRT5*, *KRT7*, *KRT8*, *KRT18*, *KRT19*, and *KRT20*). Cells with high levels of cytokeratin protein and RNA expression were defined as malignant anchors. Anchor cells were further filtered by removing outliers not near K-means cluster centroids. The averaged gene profiles of anchor cells were used as references for Insitutype to classify all cells into malignant and non-malignant categories. Batch correction across tissue slides was performed on all non-malignant cells using *ComBat* (*sva*, *v3.46.0*)^67^. Corrected non-malignant anchor cells were classified into cell types using reference profiles from a snRNA-seq dataset^23^ with each profile representing the average gene expression for that cell type. A SMI-derived reference profile was established on those anchor cells by averaging gene expression within each cell type, which was subsequently used for assigning all non-malignant cells into known cell types. Clustered cells were displayed using *Giotto* (*v2.0.0*) runUMAP function.

### 6K-SMI cell-cell interaction analysis

Ligand-receptor interactions between malignant and nerve cells were inferred using Spatially-Constrained Optimal Transport Interaction Analysis (SCOTIA v1.0.0)^24^ with default parameters. Known secreted ligand-receptor pairs were extracted from the CellChat database. Statistical significance was assessed using permutation testing, where gene expression values were randomly shuffled within each cell type to generate a null distribution. P-values were calculated based on 100 permutation iterations, with interactions considered significant at P < 0.05.

### Multiplex immunofluorescence

Candidate formalin-fixed paraffin-embedded (FFPE) human PDAC blocks that contained regions with cancer cells and PNI were sectioned (5µm) and stained as follows: Hoechst (Invitrogen, H3570), Neurofilament (ThermoFisher Scientific, Z2091MS), PanCytokeratin (eBioscience, 53-9003-82), and Pdgfd (ProteinTech, 14075-1-AP). Secondary antibodies included Goat anti-Mouse Alexa Fluor 555 (Thermo Fisher Scientific, A21127) and Donkey anti-Rabbit Alexa Fluor 647 (Thermo Fisher Scientific, A31573). Whole slide images of stained tissues were acquired using a Nikon AXR confocal at 10X magnification. Image analysis was completed using a custom workflow in QuPath in which positive cell detection was segmented on DAPI, and subsequent thresholding was performed using object classification on PanCK and Pdgfd. Nerve+ regions (< 300 µm from the nearest nerve) and Nerve-regions (> 300 µm from the nearest nerve) were identified within and across patient samples. Marker expression in relevant cell types was then quantified and compared within each patient and plotted across patients. For nerve subtype analysis, whole tissue sections and TMAs were stained with Hoechst (Invitrogen, H3570), Na_v_1.8 (Alomone Labs, ASC-016), and TH (SC25269). Na_v_1.8^+^ and TH^+^ axons were individually segmented in QuPath and quantified across patients. Axonal subtype per nerve area was calculated across individual nerves and averaged at the patient level.

### Malignant organoid derivation

The pancreas was removed from a *Kras*^LSL-G12D/+^;*Trp53*^FL/FL^;R26-LSL-dCas9-VPR-mNeonGreen female mouse at 6 weeks of age. After mincing with a razor blade, the pancreas was resuspended in 1 mL of digestion buffer (1x collagenase IV in PBS, Worthington Biochemical, LS004188) and incubated in a 37°C rotating hybridization oven in a tube for 20-30 minutes. Inside a laminar flow hood, the digested tissue was passed through a 70 μm filter over a 50mL conical and washed with 40-50 mL of PBS. After spinning for 1 minute at 800G with an acceleration of 9 and deceleration of 3, the supernatant was aspirated and the pellet was resuspended in 1 mL of PBS and moved to a 15 mL conical. After spinning for 1 minute at 800G with an acceleration and deceleration of 9, the supernatant was removed and the pellet was resuspended in a mixture of 70% Matrigel + 30% complete organoid media^68^. The cell suspension is then plated as seven 30 µL plugs in a 6-well plate, allowed to solidify in a 37°C incubator for 15 minutes, and covered in OPAC culture media.

Organoids were passaged every 4-5 days, involving manual dissociation of the Matrigel plugs and a 30-45 minute incubation in TrypLE Express Enzyme (Gibco, 12604013) at 37°C. The pellet was then replated at a 1:6-1:8 ratio. After a couple passages, the organoids were spinfected with Ad5CMVCre (University of Iowa, Ad5CMVCre) to activate the Kras mutation, Trp53 deletion, and dCas9-VPR expression. In a 24-well plate, 90k dissociated cells and 3µL of the Ad5CMVCre are combined with 500 µL of media, spun at 600G for 60 minutes at 23°C, incubated for six hours at 37°C, and then replated in three 30 µL Matrigel plugs in a 12-well plate with 1.5 mL of media. Selection, Nutlin-3a in this case, was added two days later to select for transduced cells. These organoids can be subsequently spinfected with lentivirus holding sgRNA constructs in a similar manner.

### Virus production

CRISPRa sgRNA sequences were designed using the Broad Institute’s CRISPick tool. We selected the SpyoCas9 (NGG) enzyme and chose the top-ranked sequence for each target gene. Reverse complement sequences were then generated and modified by appending “CACCG” to the 5′ end of the guide sequence, “AAAC” to the 5′ end of the reverse complement, and a terminal “C” at the 3′ end of the reverse complement. Primer pairs were ordered from IDT. Guide sequences were cloned into the lentiGuide-Puro vector (Addgene #52963) following the recommended cloning protocol. Additional packaging components, VSVG (Addgene #8454) and PsPAX2 (Addgene #12260), were prepared via standard miniprep.

HEK293T cells (Takara Bio, 632180) were seeded in 15 cm dishes to reach approximately 70% confluency the following day. For each transfection, 2 mL of Opti-MEM (Gibco, 31985062) was combined with 10 µg of guide-containing vector, 7.5 µg of PsPAX2, 2.5 µg of VSVG, and 60 µL of TransIT-LT1 (Mirus, MIR 2306). After a 30-minute incubation at room temperature, the mixture was gently pipetted onto HEK293T cells in 20 mL of fresh DMEM + 10% FBS. Following a 72-hour incubation at 37°C, viral supernatants were collected, filtered using a 0.45 µm syringe filter, and concentrated to 1.6 mL using 10 kDa Amicon Ultra-15 centrifugal filter units (Millipore Sigma, UFC901024). Concentrated virus was aliquoted and stored at –80°C until use.

### Spinfection of organoids

Organoids grown in Matrigel domes for 4–5 days were manually dissociated using a P1000 pipette and transferred to 15 mL conical tubes containing PBS. Tubes were centrifuged at 800 × g for 1 minute, and the resulting pellet was resuspended in 2–3 mL of TrypLE Express Enzyme. The tubes were incubated at 37°C, with pipetting every 10 minutes for 30–40 minutes, to promote dissociation into single cells. Once dissociated, PBS was added to fill the tube, followed by centrifugation. The cell pellet was resuspended in culture media and counted using a hemocytometer.

A total of 75,000 cells were seeded per well in a 24-well plate (GenClone, 25-107). Virus was added to each well, and culture media was added to bring the total volume to 500 µL. 3 µL of Ad-Cre (University of Iowa, Ad5CMVCre) and 100 µL of concentrated sgRNA lentivirus were used. Plates were centrifuged at 600 × g for 1 hour at 23°C, then incubated for 6 hours at 37°C. Cells were then embedded into seven 30 µL Matrigel domes and replated into 6-well plates (GenClone, 25-105). Selection agents such as nutlin and puromycin were added to the media two days post-spinoculation.

### Derivation of 2D malignant cells from 3D organoids

Malignant cells were transitioned from 3D organoid cultures to 2D adherent monolayers through serial passaging under gradually modified culture conditions. Briefly, 3D organoids were dissociated into single cells using mechanical trituration and enzymatic digestion and plated onto standard tissue culture plates in a 1:1 mixture of OPAC and DMEM 10% FBS. Cells were maintained until reaching approximately 80% confluence before passaging. At each subsequent passage, the proportion of DMEM 10% FBS was increased in a stepwise manner to 25:75 (OPAC:DMEM 10% FBS), followed by complete replacement with DMEM 10% FBS. In general, cells underwent at least four passages before displaying stable 2D morphology and growth characteristics.

### DRG harvest and dissociation

Postnatal pups (day 5-7) of the C57BL/6J background and Na_v_1.8-Cre tdTomato mice were used, and DRGs were harvested as previously described^69^ with minor modifications. Briefly, pups were sacrificed through isoflurane overdose and the spine was isolated. A longitudinal cut was made along the back of the spine to reveal the spinal cord and DRGs. The DRGs were carefully plucked and placed in 1.7 mL tubes filled with Leibovitz’s L-15 Medium (Thermo Fisher Scientific, 11415064) with 30-40 DRGs per tube. To dissociate, the tubes were spun at 800G for 5 minutes and the pellet was resuspended in 1 mL of 2mg/mL Collagenase/Dispase (Sigma-Aldrich, 11097113001). The DRGs were agitated with a closed glass Pasteur pipette for 30 seconds, and then placed on a thermomixer set to 37°C, 1000 RPM for 40 minutes. The tubes were spun down at 800G for 5 minutes, resuspended in 1 mL of TrypLE Express, and returned to the thermomixer for 15 minutes. The DRGs were then washed three times in DMEM+10% FBS and plated in pre-coated plates treated with Poly D-lysine and Laminin (Corning, 354596).

### Flow cytometry

Primary DRG cells were dissociated as described above. Cells growing in culture were dissociated using 0.05% Trypsin EDTA and washed in PBS. Single cell suspensions were pelleted at 2000 rpm, resuspended in FACS buffer (PBS with 2% BSA) with antibodies, and stained on ice for 20 minutes in the dark before being washed in PBS and resuspended in FACS buffer. Glial and cancer cells were stained with CD16/32 (Trustain FcX Plus, BioLegend, 156604) and Pdgfrb (Invitrogen, 16-1402-82) at a dilution of 1:200. Flow cytometry was performed on the Attune NxT Flow Cytometer.

### DRG neurite outgrowth assay

Dissociated neurons were plated in base media composed of 100 mL Neurobasal Plus Medium (Thermofisher, A3582901), 2 mL B-27 Plus Supplement (Thermofisher, A3582801), 250 μL GlutaMAX, 1x Penicillin-Streptomycin (Cytiva, SV30010), 5 μg FGF (Preprotech, 100-26) filtered through a 0.45 µm filter. 10,000 neurons were plated per well of a 96-well plate with 100 µL of media. After allowing the neurons to adhere overnight, the debris was gently rinsed off and the corresponding media conditions were added. Whole DRGs were plated in 7.5 µL of 100% Matrigel and cultured in DMEM with 10%FBS. Recombinant Pdgfd and/or SU16f were added to the base media to reach target concentrations. For dissociated neurons, neuronal cell body clusters and fluorescent neurites were segmented and clustered using Incucyte NeuroTrack Analysis Software Module (Sartorius, 9600-0010). For whole DRGs, axonal area was calculated as the difference between initial and final DRG fluorescence area. The net change in neurite length was measured as a function of time in relation to the first measurement and normalized to the number of cell body clusters.

### Whole DRG invasion assay

Lumbar DRGs were harvested from postnatal day 21 pups and trimmed before being placed into the center of the well in a black-bottom 96-well plate. 7.5µL of 100% Matrigel was added onto the DRG to enclose it into a dome. DRG-containing domes were placed into the incubator for 15 minutes at 37°C to solidify the Matrigel, after which 100 µL of prewarmed DMEM 10% FBS containing 12,000 cancer cells with or without recombinant Pdgfd protein (R&D Systems, 9738-SB-050) and SU16f (MCE, HY-108628) was added into the well. The neuroinvasion index was calculated by dividing the number of cancer cell spheroids invading onto the DRG by the total number of cancer spheroids. Distance was determined as the total distance traveled by each invading cancer cell spheroid. Speed was calculated by dividing the distance traveled by each individual cancer cell spheroid by the total invasion time.

### Transwell invasion assays

Pre-coated growth factor-reduced Matrigel transwell inserts with 8um polyethylene terephthalate membranes (Corning, 354483) were used for transwell invasion assays. Transwell inserts were prepared per the manufacturer’s instructions and added to a companion 24-well plate (Falcon, 353504). After serum starvation for 24 hours, 5 x 10^4^ cancer cells in 500 µL of serum-free DMEM culture media were added to the upper transwell membrane insert. In Pdgfd recombinant protein treated conditions, 20 nM of exogenous Pdgfd protein was added to the 500 µL cell suspension in serum-free DMEM media. 750 µL of DMEM + 10% FBS was added to the lower chamber wells of the companion plate. In Pdgfd recombinant protein treated conditions, 20 nM of exogenous Pdgfd protein was added to the 750 µL of DMEM + 10% FBS culture media. After culturing for 48 hours (5% CO2 at 37°C), transwell inserts were taken out of their companion plates, non-invading cancer cells were removed from the upper surface of the inserts using cotton swabs, and inserts were fixed and stained using the Differential Quik III Stain Kit (Electron Microscopy Sciences, 26096-25) per the manufacturer’s instructions. Five FOVs from each insert were imaged at 10x magnification. The average number of cells that invaded per FOV, per insert, was quantified using a custom ImageJ Macro Plugin.

### Radial invasion assay

Cancer cells (25,000 cells in 4.5 µL of 70% Matrigel) were seeded as a droplet in the middle of a well in a black-walled 96-well plate (Corning, 3904). After the Matrigel was allowed to solidify for 15 minutes at 37°C, prewarmed DMEM 10% FBS cell culture media was added to the droplet and then removed to pre-wet the bottom of the well. Immediately after, 18 µL of 100 percent Matrigel was added to the bottom of the well to enclose the initial droplet. The plate was allowed to solidify for another 15 minutes at 37°C, after which prewarmed cell culture media alone or containing recombinant Pdgfd protein with or without SU16f was added into the well. The invasion area was determined by calculating the difference in total and starting phase confluence percentage over time.

### Orthotopic transplantation

Cancer organoid lines were plated in 30 μl Matrigel (70%) domes and cultured until confluency. On the day of implantation, the cells were digested in TrypLE Express Enzyme for 40 minutes at 37°C. Resulting dissociated single cells were resuspended in 50% Matrigel and 50% OPAC and kept on ice. Orthotopic implantation of pancreatic organoids was carried out with minor adaptations from previously established protocols^70^. Dissociated organoids (100,000 cells in 100 μL 50% Matrigel/50% OPAC) were injected into the body of the pancreas of syngeneic C57BL/6J mice aged 8–12 weeks. Proper injection was confirmed by the appearance of a fluid-filled bleb within the pancreatic parenchyma, with no leakage detected. The muscle and skin were individually approximated and closed using absorbable and non-absorbable sutures, respectively. To prevent premature Matrigel polymerization, organoid mixtures were maintained on ice throughout the procedure. All animals received Buprenorphine C III (Patterson Veterinary Supply, 07-894-9214) as pre-operative analgesia and were monitored post-operatively for signs of pain or complications. Five weeks after tumor implantation, pancreases were harvested, weighed dry, fixed in 4% paraformaldehyde (PFA), placed in 30% sucrose, and embedded in OCT. 20 μm sections were stained and imaged using a Nikon AXR at 10x magnification.

### Live cell imaging

For whole DRG invasion, radial invasion, and neurite outgrowth assays, phase contrast and fluorescence images were captured every 6 hours using the Incucyte SX5 (Sartorius). Live confocal imaging was performed on a CellVoyager CQ1 Benchtop spinning disk confocal system (Yokogawa). Whole DRGs from Plp-EGFP mice and KPT cancer cells were seeded and allowed to adhere overnight before image acquisition. Images were taken every 25 minutes for approximately 24 hours. Individual channels were merged in ImageJ for analysis. Cancer cell pulling was quantified by determining the total distance traveled by individual KPT spheroids while in physical contact with EGFP+ glia. For 3D phase holotomography, samples were similarly prepared and imaged on a Tomocube HT-X1 Plus.

### Sciatic nerve invasion assay

Sciatic nerve injections were carried out in the manner as previously described^8^. Cancer cells were pretreated for 3 days with 200 µM SU16f or DMSO vehicle as a control. Mice were anesthetized using isoflurane (2-4%), and the left upper leg and flank region were shaved to remove fur. The surgical site was sterilized, and a ∼2 cm incision was made in the skin of the left upper thigh area and the muscle teased apart with surgical scissors to expose the sciatic nerve. The sciatic nerve was elevated with a stainless-steel spatula, and 15,000 cancer cells resuspended in 2 μL of media were injected into the sciatic nerve using a 10μL Hamilton syringe with a 33-gauge needle. The muscle and skin were repositioned and the skin was approximated and sutured. All animals received Meloxicam (Patterson Veterinary Supply, 078937566) as pre-operative analgesia and were monitored post-operatively for signs of pain or complications. Mice received intraperitoneal injections of SU16f (10 mg/kg) or DMSO control daily for 14 days following surgery. Sciatic nerves and tumors were harvested on day 14, fixed in 4% PFA, subsequently cleared using Methyl Salicylate as previously described^71^, and imaged whole by Nikon AXR confocal.

#### Acknowledgements

We are grateful to the patients and their families who contributed their time and surgical specimens to this study. We thank Dianne Moschella, Tina Balducci, Sharon McSorley, Serena Sullivan, Misha Pivovarov, Shannon Hoyt, Zeina Chaptini, Jen Wang, Richard Bouley, D.J. Shin, Rachel Strauss, Karen Yee, Katherine Anderson, Judy Teixeira, Margaret Magendantz, Kim Mercer, William Rideout, and Cristin McCabe for administrative and technical support. We also thank our colleagues from the animal facility. Graphics of experimental setups were created with <u>BioRender.com</u>.

## Author contributions

P.L.W. and W.L.H. developed the study concept. P.L.W., W.L.H., N.A.L., J.S., E.N.P., and T.J. designed the experiments. W.L.H., J.L.B., N.A.L. and M.M-K. designed the custom tissue microarrays and collated clinical data. M.M-K., N.C., M.G., N.A.L., and K.C. led the histological sectioning. J.S., J.A.G., H.I.H., S.H., Y.K., T.K., J.M.B., W.L.H., P.D., S.H., J.W.R., A.B., M.P., and E.M performed spatial profiling. D.G., C.S., J.C., H.C., P.L.W., J.W.R., M.H., and W.L.H. analyzed the spatial data. J.S., J.A.G., J.W.B., and D.G. designed and engineered CRISPR cell lines. P.L.W., N.A.L., E.N.P., J.S., J. H-G., A.L., P.N., and W.L.H. performed and analyzed in vitro assays. J.S., N.A.L., W.L.H., P.L.W., and G.C.J. performed and analyzed in vivo orthotopic transplants. P.L.W., D.G., C.S., J.C., E.N.P., N.A.L., W.L.H., and J.S. generated the tables and figures. T.H., J.W., H.R.. and C.F.C. supported clinical workflows. R.W., D.T.T., T.J., and L.Z. provided important scientific insights. W.L.H., M.H., T.H., D.T.T., and T.J. provided funding. W.L.H. and M.H. supervised the work. P.L.W., N.A.L., E.N.P., and W.L.H. wrote the manuscript, and all authors reviewed the manuscript.

---

**Figure S1.**
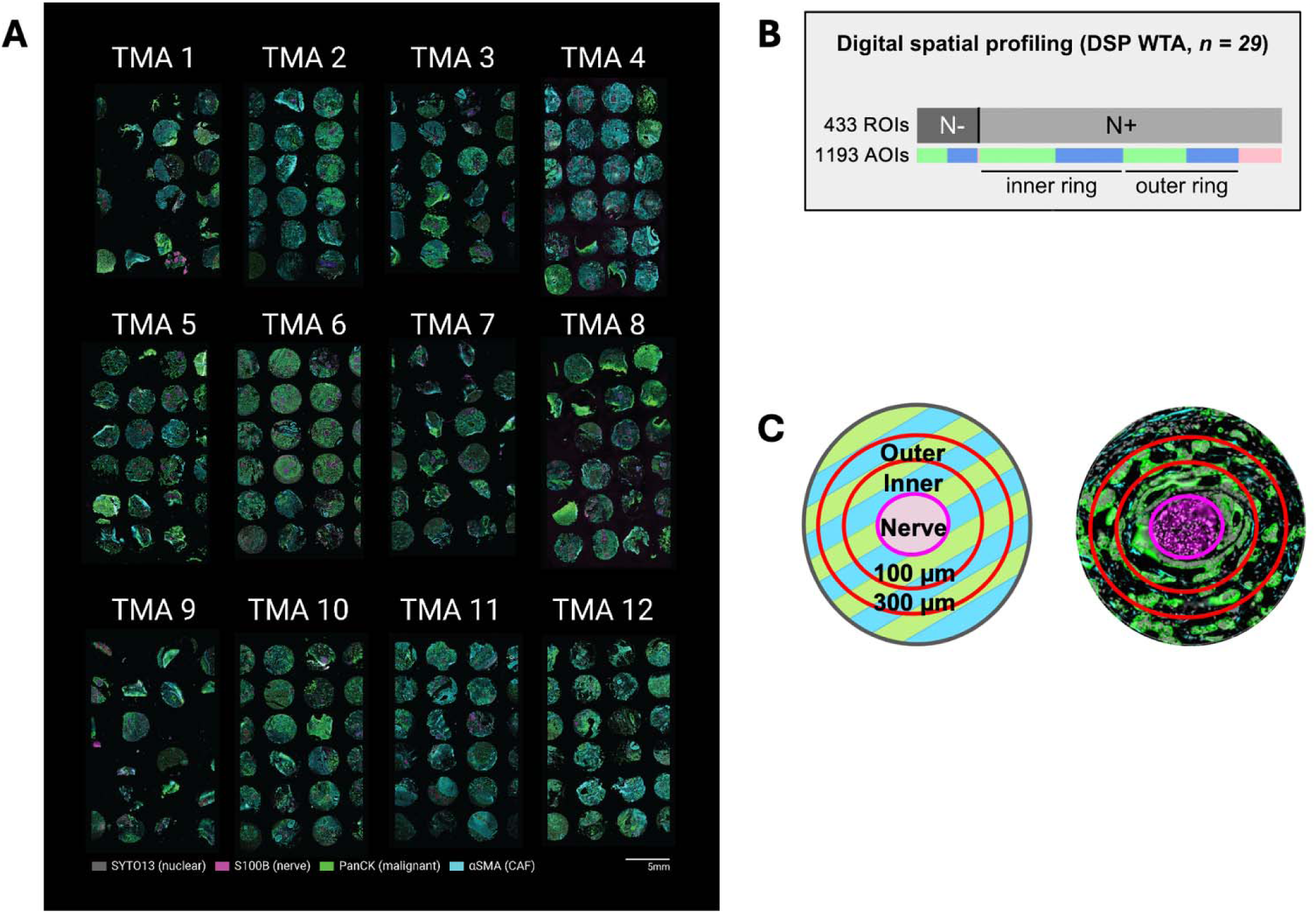
DSP experimental design. (**A**) Twelve custom human PDAC TMAs containing N+ and N-cores for WT-DSP and 6K SMI visualized with SYTO13 (gray) nuclear stain, S100B (magenta) nerve stain, PanCK (green) malignant stain, and LSMA (cyan) CAF stain. (**B**) WT-DSP study design for unbiased whole-transcriptome screening performed on 433 ROIs across 29 patients, separated into 1193 cell type-and nerve proximity-specific AOIs. ROIs are separated into nerve-positive (N+, n = 260) and nerve-negative (N-, n = 173) cores. AOIs are separated into epithelial (colored green, n = 170 N-, n = 231 N+ in inner ring, n = 179 N+ in outer ring), cancer-associated fibroblasts (CAFs, colored blue, n = 161 N-, n = 192 N+ in inner ring, n = 137 N+ in outer ring), and nerve (colored magenta, n = 123 N+) segments. (**C**) Schematic of concentric ROIs within N+ core.

**Figure S2.**
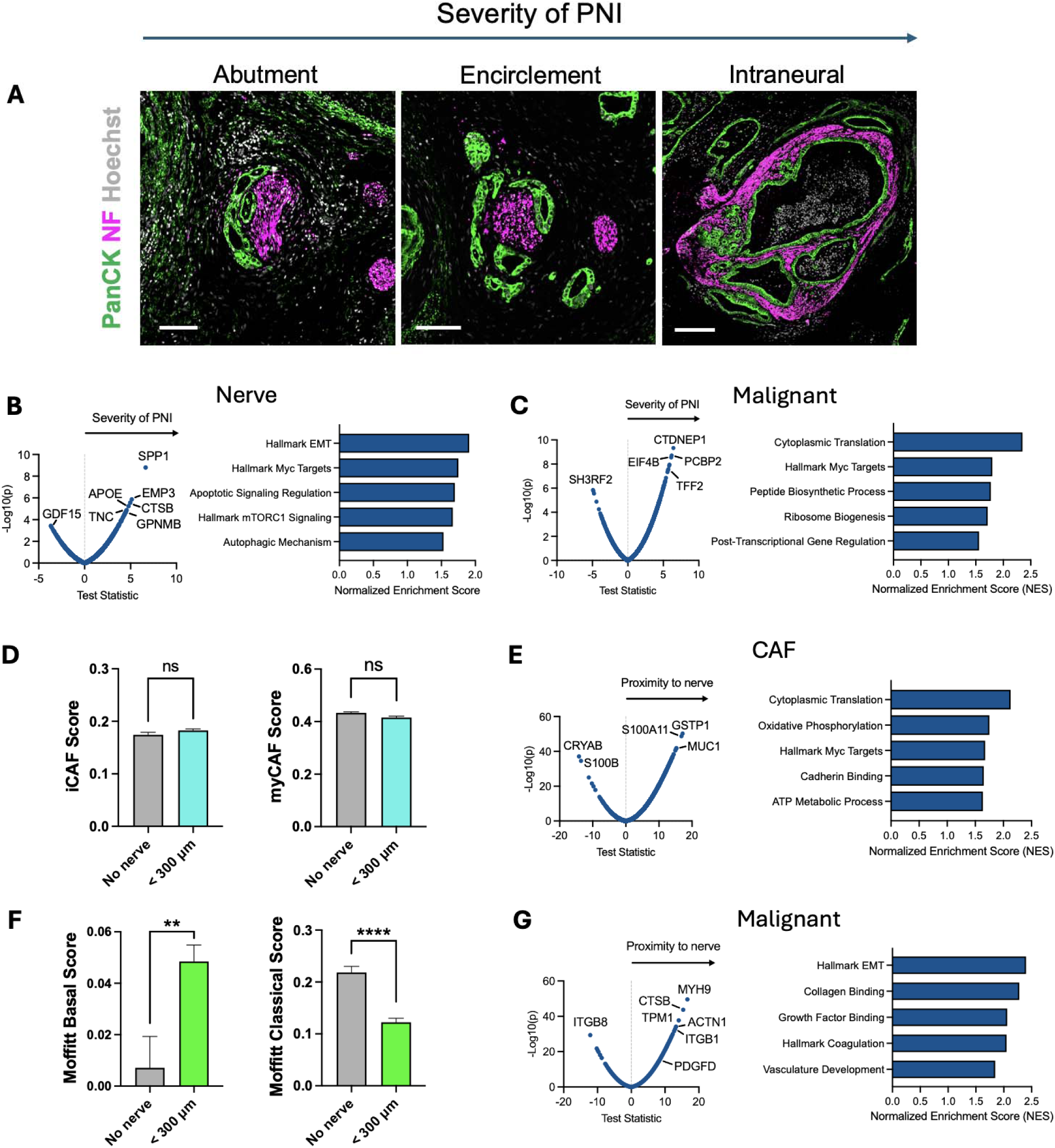
Landmark-based DSP analysis. (**A**) Representative images of DSP ROIs showing increasing severity of PNI based on the degree of nerve contact/involvement. Abutment and encirclement scale bar, 100 µm. Intraneural scale bar, 200 µm. (**B**) Genes enriched in nerves and corresponding GSEA analysis (right) as a function of increasing PNI severity. (**C**) Genes enriched in malignant cells and corresponding GSEA analysis (right) as a function of increasing PNI severity. (**D**) Normalized CAF program scores for epithelial AOIs in N-versus all N+ ROIs. iCAF (left) and myCAF (right) scores in CAFs based on nerve proximity. (**E**) Genes enriched in CAFs (left) and corresponding GSEA analysis (right) as a function of increasing proximity to nerves. (**F**) Normalized malignant program scores for epithelial AOIs in N-versus all N+ ROIs. Moffitt Basal (left) and Moffitt Classical (right) scores in malignant cells based on nerve proximity. *\*\*P* < 0.01, *\*\*\*\*P* < 0.0001 (Mann-Whitney test, Mean SEM). (**G**) Genes enriched in malignant cells (left) and corresponding GSEA analysis (right) as a function of increasing proximity to nerves.

**Figure S3.**
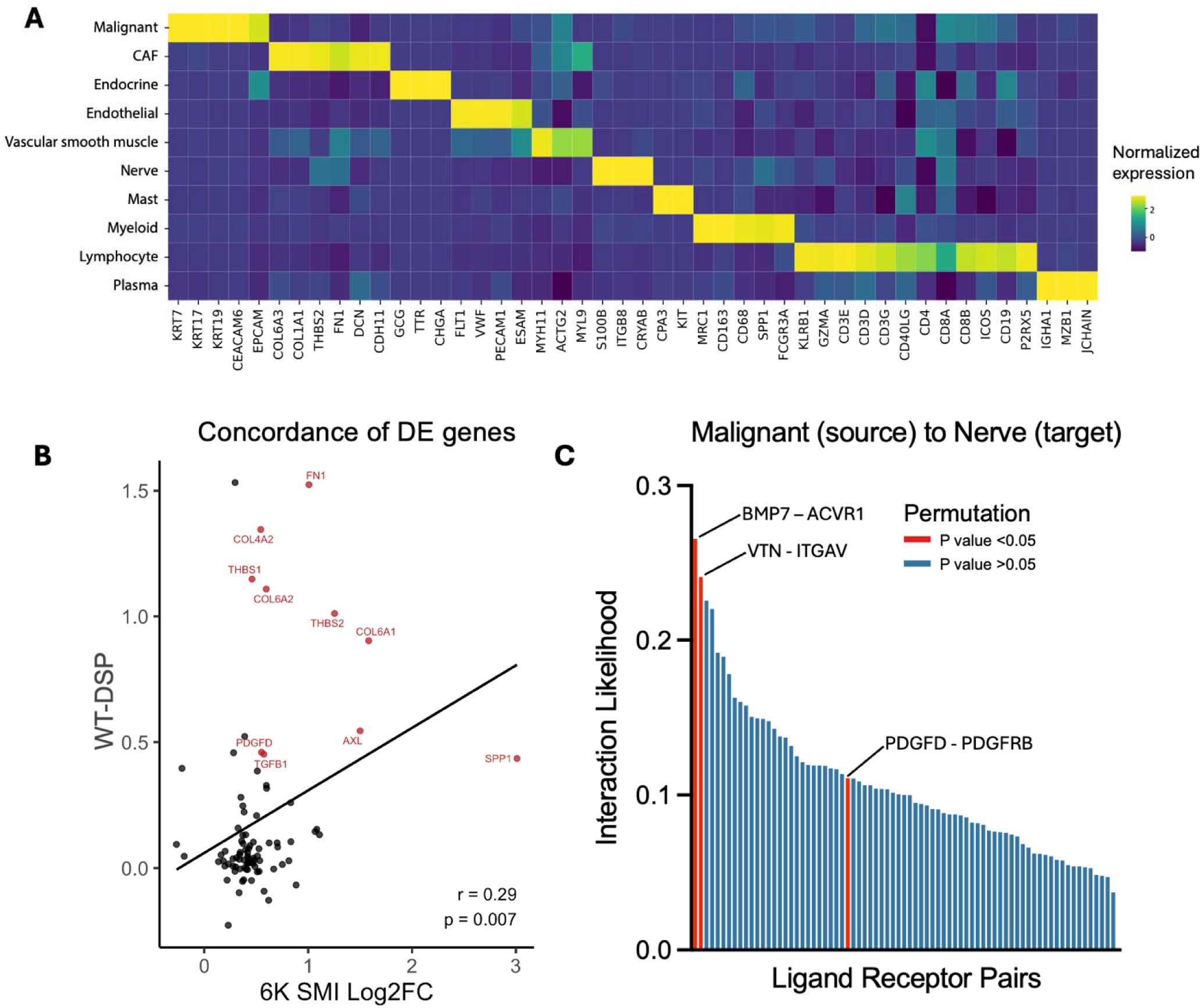
Single cell SMI and concordance analysis. (**A**) Heat map of 6K-SMI showing expression of select marker genes for annotated cell types. (**B**) Concordance of DE genes from PNI+ malignant cells between WT-DSP and 6K-SMI. (**C**) Interaction likelihood score of LR interactions between malignant ligands and nerve receptors. LR pairs with P value < 0.05 are highlighted in red.

**Figure S4.**
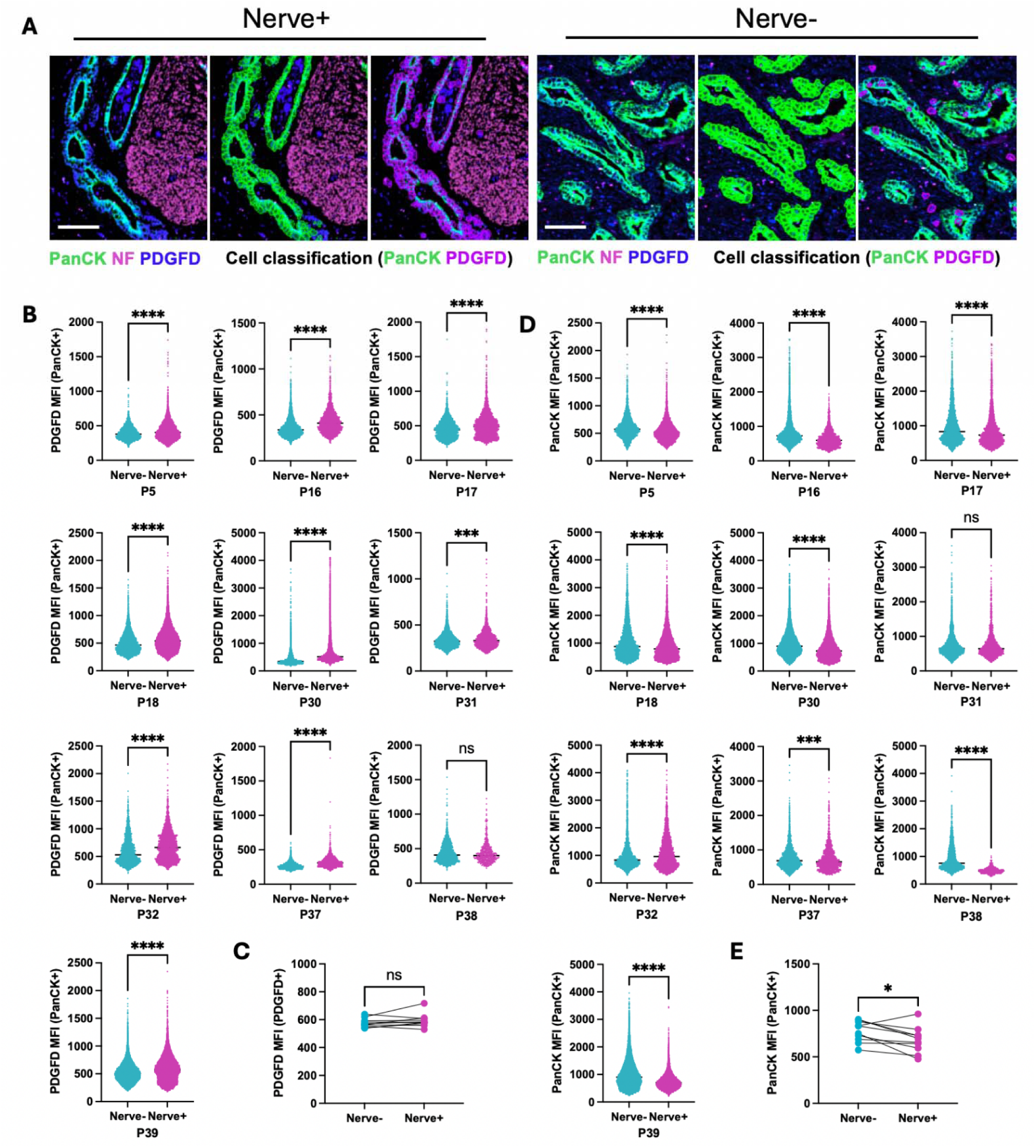
Quantitative IF assessment of PDGFD expression. (**A**) IF images of an example N+ (left) and N-(right) field of view of human PDAC with cell objects of interest, including PanCK+ malignant cells (green) and PDGFD+ cells (purple) defined using QuPath for quantitative IF analysis. Scale bars, 100 µm. (**B**) Plots of PDGFD MFI expression values from individual PDAC patients corresponding to **Fig. 1I** in Nerve-(> 300 µm from the nearest nerve) and Nerve+ (< 300 µm from the nearest nerve) regions (each dot corresponds to one cell). (**C**) Mean values of PDGFD MFI in PanCK-PDGFD+ cells (each dot corresponds to a single patient). (**D**) Plots of individual PanCK expression values (each dot corresponds to one cell) between Nerve+ (< 300 um) and Nerve-regions corresponding to panel e. (**E**) Mean values of PanCK MFI in PanCK+ PDGFD+ cells (each dot corresponds to a single patient). *\*P* < 0.05, \*\*\**P* < 0.001, *\*\*\*\*P* < 0.0001 [Mann-Whitney test, Mean SD (B-D); Wilcoxon test (E)].

**Figure S5.**
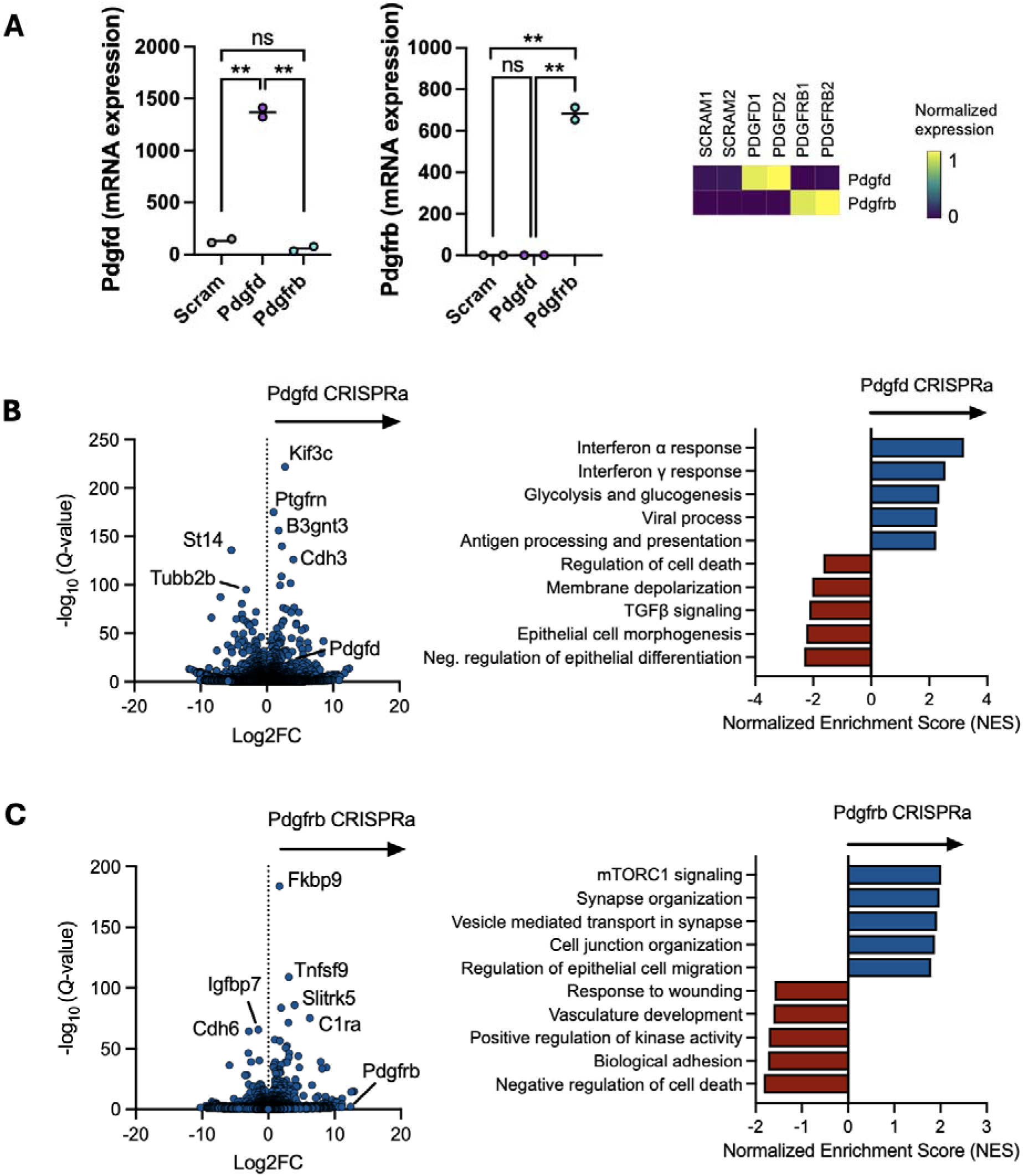
Transcriptomic analysis of CRISPRa lines. (**A**) Pdgfd (left) and Pdgfrb (middle) mRNA expression from bulk RNAseq of CRISPRa and scramble gRNA control lines (n = 2). *\*\*P* < 0.01 (Unpaired t test). Heat map representation of CRISPRa and scramble control line expression of Pdgfd and Pdgfrb. (**B**) Volcano plot of genes enriched in Pdgfd-overexpressing cell line (left) and corresponding GSEA analysis (right). (**C**) Volcano plot of genes enriched in Pdgfrb-overexpressing cell line (left) and corresponding GSEA analysis (right).

**Figure S6.**
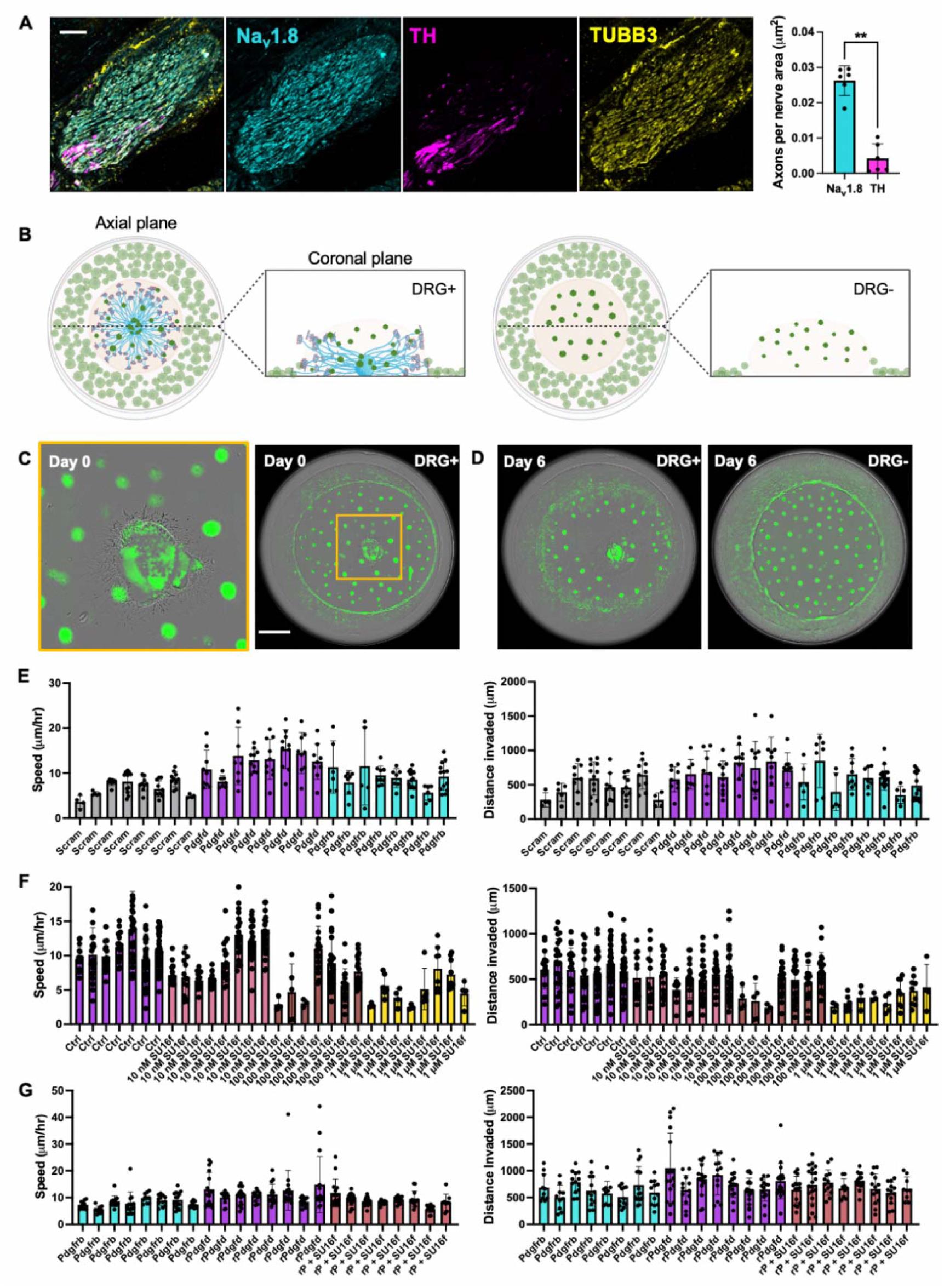
Rationale and schematic for whole DRG invasion assay. (**A**) Nerve subtype staining (left) and quantification (right) for Na_v_1.8 (cyan) and TH (magenta) in human PDAC (n = 6 patients). *\*\*P* < 0.01 (Mann-Whitney test, Mean SD). (**B**) Schematic of whole DRG invasion assay. **(C)** Still frame from live cell imaging of central DRG explant and whole DRG invasion assay at day 0. Cancer cells are visualized by mNeon reen expression. Scale bar, 1 mm. (**D**) End point (day 6) image of whole DRG invasion from (b, left) and DRG-negative control (right). (**E**) Compiled replicates for speed and distance analyses of Pdgfd, Pdgfrb, and control cancer cell invasion of DRGs over 6 days. (**F**) Compiled replicates for speed and distance analyses of Pdgfd cancer cell invasion of DRGs with or without SU16f treatment over 6 days. (**G**) Compiled replicates for speed and distance analyses of Pdgfrb cancer cell invasion of DRGs with or without exogenous recombinant Pdgfd over 6 days. Abbreviations: Scram = Scramble, rP/rPdgfd = recombinant Pdgfd.

**Figure S7.**
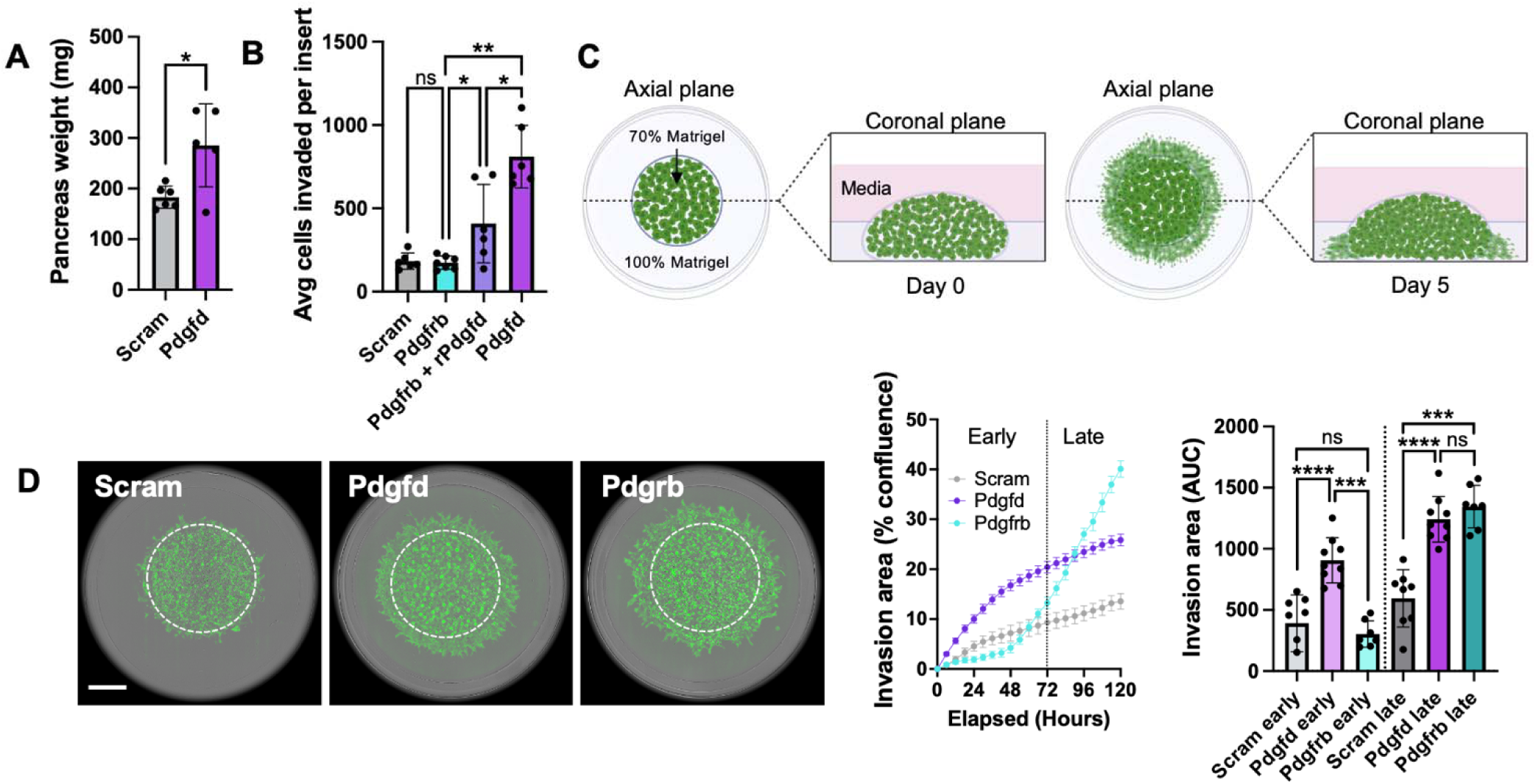
Radial invasion assay. (**A**) Pancreas weight from mice 5 weeks after receiving Pdgfd-overexpressing and control orthotopic tumors. Corresponds to **Fig. 3b** (n = 5 mice). (**B**) Transwell invasion assay with Scramble control, Pdgfrb line with or without recombinant 20 nM Pdgfd, and Pdgfd line (n = 6). (**C**) Schematic of radial invasion assay. (**D**) Incucyte images (day 4) and graph of invasion area over time for Scramble, Pdgfd, and Pdgfrb lines (left and middle). Scale bar, 1 mm. Quantification of AUC for invasion area from 0-72 hours and from 72-120 hours (n = 7-9). *\*P* < 0.05, *\*\*P* < 0.01, \*\*\**P* < 0.001, *\*\*\*\*P* < 0.0001 [Unpaired t test, Mean SD (A); Mann-Whitney test, Mean SD (B), (D)]. Abbreviations: Scram = Scramble, rPdgfd = recombinant Pdgfd.

**Fig. S8.**
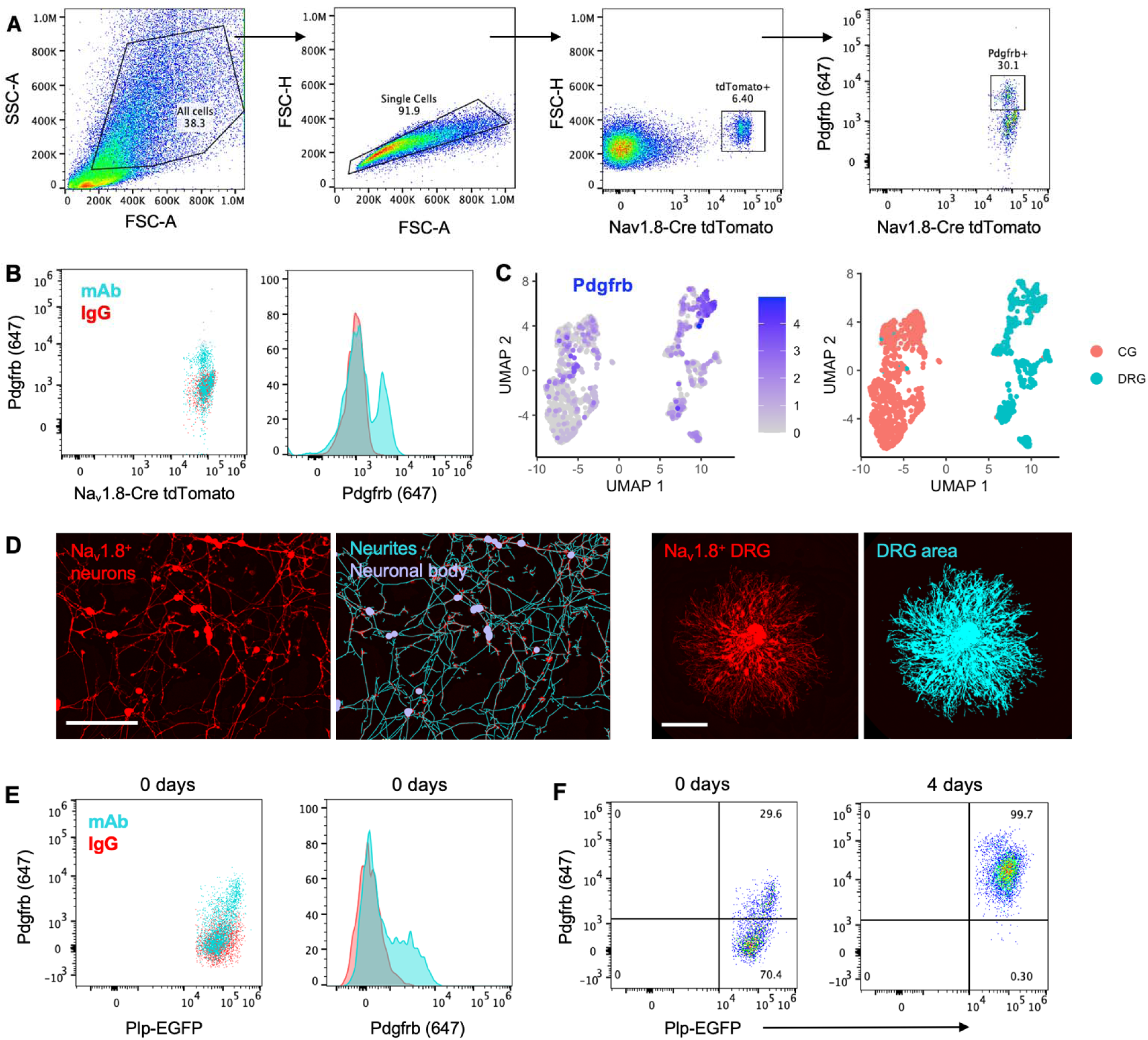
Neuronal Pdgfrb expression and neurite tracking. (**A**) Gating strategy for determining Pdgfrb expression in tdTomato+ neurons from Na_v_1.8Cre-tdTomato mice. (**B**) Flow cytometry plots showing Pdgfrb expression in freshly isolated tdTomato+ neurons from Na_v_1.8-Cre-tdTomato mice. Blue plots correspond to Pdgfrb-stained and red plots correspond to IgG isotype control-stained condition. (**C**) UMAP plots showing Pdgfrb expression in celiac ganglia (CG) and dorsal root ganglia (DRG) innervating normal pancreas and pancreatic tumors in mice adapted from Thiel et al. (*36*). (**D**) Sample images of Na_v_1.8-Cre tdTomato+ dissociated nerves taken with live cell imaging (Incucyte SX5) with tracing of neurites (cyan) and neuronal cell bodies (purple) at 48 hours by NeuroTrack analysis module for neurite outgrowth quantification (left). Scale bar, 200 µm. hole tdTomato+ (red) DRG and red object mask (cyan) of DRG outgrowth area at 120 hours (right). Scale bar, 1 mm. (**E**) Flow cytometry plot and histogram showing Pdgfrb expression in freshly dissociated EGFP+ glial cells from Plp-EGFP mice. Blue plots correspond to Pdgfrb-stained and red plots correspond to IgG-stained condition. (**F**) Flow cytometry plots showing the percentage of EGFP+ glia expressing Pdgfrb 0 days versus 4 days after culturing.

**Table 1.**
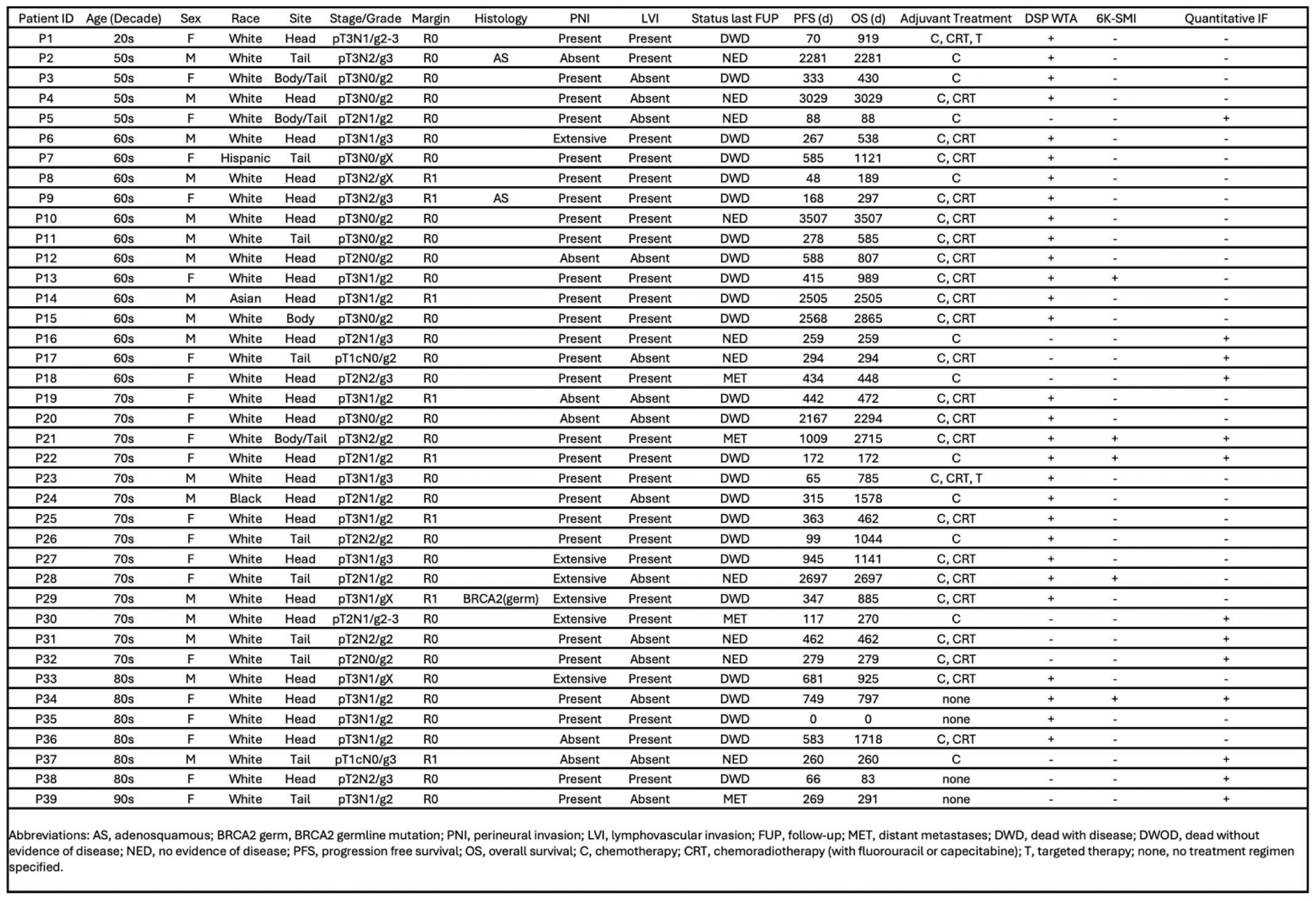
Patient cohort and clinicopathologic data.

## References

1. Shi DD, Guo JA, Hoffman HI, et al. Therapeutic avenues for cancer neuroscience: translational frontiers and clinical opportunities. Lancet Oncol. 2022;23(2):e62–e74. doi:10.1016/S1470-2045(21)00596-9

2. Liebig C, Ayala G, Wilks JA, Berger DH, Albo D. Perineural invasion in cancer: a review of the literature. Cancer. 2009;115(15):3379–3391. doi:10.1002/cncr.24396

3. Chatterjee D, Katz MH, Rashid A, et al. Perineural and intraneural invasion in posttherapy pancreaticoduodenectomy specimens predicts poor prognosis in patients with pancreatic ductal adenocarcinoma. Am J Surg Pathol. 2012;36(3):409–417. doi:10.1097/PAS.0b013e31824104c5

4. Demir IE, Ceyhan GO, Liebl F, D’Haese JG, Maak M, Friess H. Neural invasion in pancreatic cancer: the past, present and future. Cancers (Basel). 2010;2(3):1513–1527. doi:10.3390/cancers2031513

5. Renz BW, Takahashi R, Tanaka T, et al. β2 Adrenergic-Neurotrophin Feedforward Loop Promotes Pancreatic Cancer. Cancer Cell. 2018;33(1):75–90.e7. doi:10.1016/j.ccell.2017.11.007

6. Biankin A V, Waddell N, Kassahn KS, et al. Pancreatic cancer genomes reveal aberrations in axon guidance pathway genes. Nature. 2012;491(7424):399–405. doi:10.1038/nature11547

7. Silverman DA, Martinez VK, Dougherty PM, Myers JN, Calin GA, Amit M. Cancer-Associated Neurogenesis and Nerve-Cancer Cross-talk. Cancer Res. 2021;81(6):1431-p1440. doi:10.1158/0008-5472.CAN-20-2793

8. Deborde S, Gusain L, Powers A, et al. Reprogrammed Schwann Cells Organize into Dynamic Tracks that Promote Pancreatic Cancer Invasion. Cancer Discov. 2022;12(10):2454–2473. doi:10.1158/2159-8290.CD-21-1690

9. Deborde S, Wong RJ. The Role of Schwann Cells in Cancer. Adv Biol. 2022;6(9):e2200089. doi:10.1002/adbi.202200089

10. Guo JA, Hoffman HI, Shroff SG, et al. Pan-cancer Transcriptomic Predictors of Perineural Invasion Improve Occult Histopathologic Detection. Clin Cancer Res. 2021;27(10):2807–2815. doi:10.1158/1078-0432.CCR-20-4382

11. Merritt CR, Ong GT, Church SE, et al. Multiplex digital spatial profiling of proteins and RNA in fixed tissue. Nat Biotechnol. 2020;38(5):586–599. doi:10.1038/s41587-020-0472-9

12. Liu X, Sun Y, Li H, et al. Effect of Spp1 on nerve degeneration and regeneration after rat sciatic nerve injury. BMC Neurosci. 2017;18(1):30. doi:10.1186/s12868-017-0348-1

13. Li S, Jakobs TC. Secreted phosphoprotein 1 slows neurodegeneration and rescues visual function in mouse models of aging and glaucoma. Cell Rep. 2022;41(13):111880. doi:10.1016/J.CELREP.2022.111880

14. Brosius Lutz A, Lucas TA, Carson GA, et al. An RNA-sequencing transcriptome of the rodent Schwann cell response to peripheral nerve injury. J Neuroinflammation. 2022;19(1):105. doi:10.1186/s12974-022-02462-6

15. Schmitd LB, Perez-Pacheco C, Bellile EL, et al. Spatial and Transcriptomic Analysis of Perineural Invasion in Oral Cancer. Clin Cancer Res. 2022;28(16):3557–3572. doi:10.1158/1078-0432.CCR-21-4543

16. Baruch EN, Nagarajan P, Gleber-Netto FO, et al. Inflammation induced by tumor-associated nerves promotes resistance to anti-PD-1 therapy in cancer patients and is targetable by interleukin-6 blockade. Res Sq. Published online July 18, 2023. doi:10.21203/rs.3.rs-3161761/v1

17. Elyada E, Bolisetty M, Laise P, et al. Cross-Species Single-Cell Analysis of Pancreatic Ductal Adenocarcinoma Reveals Antigen-Presenting Cancer-Associated Fibroblasts. Cancer Discov. 2019;9(8):1102–1123. doi:10.1158/2159-8290.CD-19-0094

18. Moffitt RA, Marayati R, Flate EL, et al. Virtual microdissection identifies distinct tumor-and stroma-specific subtypes of pancreatic ductal adenocarcinoma. Nat Genet. 2015;47(10):1168–1178. doi:10.1038/ng.3398

19. Amit M, Na’ara S, Gil Z. Mechanisms of cancer dissemination along nerves. Nat Rev Cancer. 2016;16(6):399–408. doi:10.1038/nrc.2016.38

20. Wu Q, Hou X, Xia J, et al. Emerging roles of PDGF-D in EMT progression during tumorigenesis. Cancer Treat Rev. 2013;39(6):640–646. doi:10.1016/j.ctrv.2012.11.006

21. Funa K, Sasahara M. The roles of PDGF in development and during neurogenesis in the normal and diseased nervous system. J Neuroimmune Pharmacol. 2014;9(2):168–181. doi:10.1007/s11481-013-9479-z

22. Pandey P, Khan F, Upadhyay TK, Seungjoon M, Park MN, Kim B. New insights about the PDGF/PDGFR signaling pathway as a promising target to develop cancer therapeutic strategies. Biomedicine & Pharmacotherapy. 2023;161:114491. doi:10.1016/J.BIOPHA.2023.114491

23. Hwang WL, Jagadeesh KA, Guo JA, et al. Single-nucleus and spatial transcriptome profiling of pancreatic cancer identifies multicellular dynamics associated with neoadjuvant treatment. Nat Genet. 2022;54(8):1178–1191. doi:10.1038/s41588-022-01134-8

24. Shiau C, Cao J, Gong D, et al. Spatially resolved analysis of pancreatic cancer identifies therapy-associated remodeling of the tumor microenvironment. Nat Genet. 2024;56(11):2466–2478. doi:10.1038/s41588-024-01890-9

25. Nassar MA, Stirling LC, Forlani G, et al. Nociceptor-specific gene deletion reveals a major role for Nav1.7 (PN1) in acute and inflammatory pain. Proc Natl Acad Sci U S A. 2004;101(34):12706–12711. doi:10.1073/pnas.0404915101

26. Zimmermann K, Leffler A, Babes A, et al. Sensory neuron sodium channel Nav1.8 is essential for pain at low temperatures. Nature. 2007;447(7146):855–858. doi:10.1038/nature05880

27. Zheng Y, Liu P, Bai L, Trimmer JS, Bean BP, Ginty DD. Deep Sequencing of Somatosensory Neurons Reveals Molecular Determinants of Intrinsic Physiological Properties. Neuron. 2019;103(4):598–616.e7. doi:10.1016/j.neuron.2019.05.039

28. Bergsten E, Uutela M, Li X, et al. PDGF-D is a specific, protease-activated ligand for the PDGF beta-receptor. Nat Cell Biol. 2001;3(5):512–516. doi:10.1038/35074588

29. Sun L, Tran N, Liang C, et al. Design, synthesis, and evaluations of substituted 3-[(3-or 4-carboxyethylpyrrol-2-yl)methylidenyl]indolin-2-ones as inhibitors of VEGF, FGF, and PDGF receptor tyrosine kinases. J Med Chem. 1999;42(25):5120–5130. doi:10.1021/jm9904295

30. Huang F, Wang M, Yang T, et al. Gastric cancer-derived MSC-secreted PDGF-DD promotes gastric cancer progression. J Cancer Res Clin Oncol. 2014;140(11):1835–1848. doi:10.1007/s00432-014-1723-2

31. Boj SF, Hwang CI, Baker LA, et al. Organoid models of human and mouse ductal pancreatic cancer. Cell. 2015;160(1-2):324–338. doi:10.1016/j.cell.2014.12.021

32. Kapałczyńska M, Kolenda T, Przybyła W, et al. 2D and 3D cell cultures - a comparison of different types of cancer cell cultures. Arch Med Sci. 2018;14(4):910–919. doi:10.5114/aoms.2016.63743

33. Puls TJ, Tan X, Husain M, Whittington CF, Fishel ML, Voytik-Harbin SL. Development of a Novel 3D Tumor-tissue Invasion Model for High-throughput, High-content Phenotypic Drug Screening. Sci Rep. 2018;8(1):13039. doi:10.1038/s41598-018-31138-6

34. Hudkins KL, Gilbertson DG, Carling M, et al. Exogenous PDGF-D is a potent mesangial cell mitogen and causes a severe mesangial proliferative glomerulopathy. J Am Soc Nephrol. 2004;15(2):286–298. doi:10.1097/01.asn.0000108522.79652.63

35. Hye Kim J, Gyu Park S, Kim WK, Song SU, Sung JH. Functional regulation of adipose-derived stem cells by PDGF-D. Stem Cells. 2015;33(2):542–556. doi:10.1002/stem.1865

36. Thiel V, Renders S, Panten J, et al. Characterization of single neurons reprogrammed by pancreatic cancer. Nature. 2025;640(8060):1042-1051. doi:10.1038/s41586-025-08735-3

37. Carr MJ, Toma JS, Johnston APW, et al. Mesenchymal Precursor Cells in Adult Nerves Contribute to Mammalian Tissue Repair and Regeneration. Cell Stem Cell. 2019;24(2):240–256.e9. doi:10.1016/j.stem.2018.10.024

38. Avraham O, Deng PY, Jones S, et al. Satellite glial cells promote regenerative growth in sensory neurons. Nat Commun. 2020;11(1):4891. doi:10.1038/s41467-020-18642-y

39. Fallon JR. Preferential outgrowth of central nervous system neurites on astrocytes and Schwann cells as compared with nonglial cells in vitro. J Cell Biol. 1985;100(1):198–207. doi:10.1083/jcb.100.1.198

40. Yim AKY, Wang PL, Bermingham JR, et al. Disentangling glial diversity in peripheral nerves at single-nuclei resolution. Nat Neurosci. 2022;25(2):238–251. doi:10.1038/s41593-021-01005-1

41. Mallon BS, Shick HE, Kidd GJ, Macklin WB. Proteolipid promoter activity distinguishes two populations of NG2-positive cells throughout neonatal cortical development. J Neurosci. 2002;22(3):876–885. doi:10.1523/JNEUROSCI.22-03-00876.2002

42. Miller MJ, Kangas CD, Macklin WB. Neuronal expression of the proteolipid protein gene in the medulla of the mouse. J Neurosci Res. 2009;87(13):2842–2853. doi:10.1002/jnr.22121

43. Harlow DE, Saul KE, Culp CM, Vesely EM, Macklin WB. Expression of proteolipid protein gene in spinal cord stem cells and early oligodendrocyte progenitor cells is dispensable for normal cell migration and myelination. J Neurosci. 2014;34(4):1333–1343. doi:10.1523/JNEUROSCI.2477-13.2014

44. Winkler F, Venkatesh HS, Amit M, et al. Cancer neuroscience: State of the field, emerging directions. Cell. 2023;186(8):1689–1707. doi:10.1016/j.cell.2023.02.002

45. Steller EJA, Raats DA, Koster J, et al. PDGFRB promotes liver metastasis formation of mesenchymal-like colorectal tumor cells. Neoplasia. 2013;15(2):204–217. doi:10.1593/neo.121726

46. Byun JS, Gardner K. Wounds that will not heal: pervasive cellular reprogramming in cancer. Am J Pathol. 2013;182(4):1055–1064. doi:10.1016/j.ajpath.2013.01.009

47. Hanahan D, Weinberg RA. Hallmarks of cancer: the next generation. Cell. 2011;144(5):646–674. doi:10.1016/j.cell.2011.02.013

48. MacCarthy-Morrogh L, Martin P. The hallmarks of cancer are also the hallmarks of wound healing. Sci Signal. 2020;13(648). doi:10.1126/scisignal.aay8690

49. Di Chiaro P, Nacci L, Arco F, et al. Mapping functional to morphological variation reveals the basis of regional extracellular matrix subversion and nerve invasion in pancreatic cancer. Cancer Cell. 2024;42(4):662–681.e10. doi:10.1016/J.CCELL.2024.02.017

50. Chen MM, Gao Q, Ning H, et al. Integrated single-cell and spatial transcriptomics uncover distinct cellular subtypes involved in neural invasion in pancreatic cancer. Cancer Cell. Published online July 17, 2025. doi:10.1016/j.ccell.2025.06.020

51. Johnston APW, Miller FD. The Contribution of Innervation to Tissue Repair and Regeneration. Cold Spring Harb Perspect Biol. 2022;14(9). doi:10.1101/cshperspect.a041233

52. Kumar A, Brockes JP. Nerve dependence in tissue, organ, and appendage regeneration. Trends Neurosci. 2012;35(11):691–699. doi:10.1016/j.tins.2012.08.003

53. Boilly B, Faulkner S, Jobling P, Hondermarck H. Nerve Dependence: From Regeneration to Cancer. Cancer Cell. 2017;31(3):342–354. doi:10.1016/j.ccell.2017.02.005

54. Gracia F, Sanchez-Laorden B, Gomez-Sanchez JA. Schwann cells in regeneration and cancer: an epithelial-mesenchymal transition perspective. Open Biol. 2025;15(3):240337. doi:10.1098/rsob.240337

55. Dai W, Miao Y, Yi S. Schwann cell reprogramming via EMT-like program following peripheral nerve injury and during nerve regeneration. Front Cell Dev Biol. 2025;13. doi:10.3389/fcell.2025.1621380

56. Johnston APW, Yuzwa SA, Carr MJ, et al. Dedifferentiated Schwann Cell Precursors Secreting Paracrine Factors Are Required for Regeneration of the Mammalian Digit Tip. Cell Stem Cell. 2016;19(4):433–448. doi:10.1016/j.stem.2016.06.002

57. Lefevere E, Van Hove I, Sergeys J, et al. PDGF as an Important Initiator for Neurite Outgrowth Associated with Fibrovascular Membranes in Proliferative Diabetic Retinopathy. Curr Eye Res. 2022;47(2):277–286. doi:10.1080/02713683.2021.1966479

58. Xue M, Zhu Y, Jiang Y, et al. Schwann cells regulate tumor cells and cancer-associated fibroblasts in the pancreatic ductal adenocarcinoma microenvironment. Nat Commun. 2023;14(1):4600. doi:10.1038/s41467-023-40314-w

59. Jackson EL, Willis N, Mercer K, et al. Analysis of lung tumor initiation and progression using conditional expression of oncogenic K-ras. Genes Dev. 2001;15(24):3243–3248. doi:10.1101/gad.943001

60. Marino S, Vooijs M, van Der Gulden H, Jonkers J, Berns A. Induction of medulloblastomas in p53-null mutant mice by somatic inactivation of Rb in the external granular layer cells of the cerebellum. Genes Dev. 2000;14(8):994–1004.

61. Freed-Pastor WA, Lambert LJ, Ely ZA, et al. The CD155/TIGIT axis promotes and maintains immune evasion in neoantigen-expressing pancreatic cancer. Cancer Cell. 2021;39(10):1342–1360.e14. doi:10.1016/j.ccell.2021.07.007

62. He S, Bhatt R, Brown C, et al. High-plex imaging of RNA and proteins at subcellular resolution in fixed tissue by spatial molecular imaging. Nat Biotechnol. 2022;40(12):1794–1806. doi:10.1038/s41587-022-01483-z

63. Stringer C, Wang T, Michaelos M, Pachitariu M. Cellpose: a generalist algorithm for cellular segmentation. Nat Methods. 2021;18(1):100–106. doi:10.1038/s41592-020-01018-x

64. Pachitariu M, Stringer C. Cellpose 2.0: how to train your own model. Nat Methods. 2022;19(12):1634–1641. doi:10.1038/s41592-022-01663-4

65. Petukhov V, Xu RJ, Soldatov RA, et al. Cell segmentation in imaging-based spatial transcriptomics. Nat Biotechnol. 2022;40(3):345–354. doi:10.1038/s41587-021-01044-w

66. Danaher P, et al. Insitutype: likelihood-based cell typing for single cell spatial transcriptomics. *bioRxiv*. Published online 2022.

67. Johnson WE, Li C, Rabinovic A. Adjusting batch effects in microarray expression data using empirical Bayes methods. Biostatistics. 2007;8(1):118–127. doi:10.1093/biostatistics/kxj037

68. Raghavan S, Winter PS, Navia AW, et al. Microenvironment drives cell state, plasticity, and drug response in pancreatic cancer. Cell. 2021;184(25):6119–6137.e26. doi:10.1016/j.cell.2021.11.017

69. Jurcak NR, Rucki AA, Muth S, et al. Axon Guidance Molecules Promote Perineural Invasion and Metastasis of Orthotopic Pancreatic Tumors in Mice. Gastroenterology. 2019;157(3):838–850.e6. doi:10.1053/j.gastro.2019.05.065

70. Kim MP, Evans DB, Wang H, Abbruzzese JL, Fleming JB, Gallick GE. Generation of orthotopic and heterotopic human pancreatic cancer xenografts in immunodeficient mice. Nat Protoc. 2009;4(11):1670–1680. doi:10.1038/nprot.2009.171

71. Randolph GJ, Bala S, Rahier JF, et al. Lymphoid Aggregates Remodel Lymphatic Collecting Vessels that Serve Mesenteric Lymph Nodes in Crohn Disease. Am J Pathol. 2016;186(12):3066–3073. doi:10.1016/j.ajpath.2016.07.026

